# The APC/C^CCS52A2^ E3-Ligase Complex targeted PKN1 Protein is a Plant Lineage-Specific Driver of Cell Division

**DOI:** 10.1101/2021.09.27.462019

**Authors:** Alex Willems, Yuanke Liang, Jefri Heyman, Thomas Eekhout, Hilde Van den Daele, Ilse Vercauteren, Lieven De Veylder

## Abstract

The anaphase-promoting complex/cyclosome (APC/C) marks key cell cycle proteins for proteasomal breakdown, thereby ensuring unidirectional progression through the cell cycle. Its target recognition is temporally regulated by activating subunits, one of which is called CELL CYCLE SWITCH 52 A2 (CCS52A2). We sought to expand the knowledge of identified APC/C targets by using the severe growth phenotypes of *CCS52A2*-deficient *Arabidopsis thaliana* plants as a readout in a suppressor mutagenesis screen, resulting in the identification of the previously undescribed gene called *PIKMIN1* (*PKN1*). PKN1 deficiency rescues the disorganized root stem cell phenotype of the *ccs52a2-1* mutant, whereas an excess of PKN1 inhibits growth of *ccs52a2-1* plants, indicating the importance of PKN1 abundance for proper development. Accordingly, the lack of PKN1 in a wild-type background negatively impacts cell division, while its ectopic expression promotes proliferation. PKN1 shows a cell cycle phase-dependent accumulation pattern, localizing to microtubular structures, including the preprophase band, the mitotic spindle, and phragmoplast. *PKN1* is conserved throughout the plant kingdom, with its function in cell division being evolutionary conserved in the liverwort *Marchantia polymorpha*. Our data thus demonstrate that PKN1 represents a novel, plant-specific gene with a rate-limiting role in cell division, which is proteolytically controlled by the CCS52A2-activated APC/C.

**One-Sentence Summary:** PKN1 is a conserved plant-specific protein that is rate-limiting for cell division, likely through its interaction with microtubuli, and is proteolytically controlled by APC/C^CCS52A2^.

## INTRODUCTION

The selective destruction of proteins is essential to provide unidirectionality and finality to cellular processes such as the cell cycle. One way to achieve this is through the ubiquitin-proteasome pathway. Ubiquitination is the process of attaching a ubiquitin moiety to a target protein, which results in its recognition and proteolytic destruction by the 26S proteasome (Glickman and Ciechanover, 2002). The process of ubiquitination involves a set of enzymatic reactions: an E1 ubiquitin-activating enzyme binds and activates the ubiquitin in an ATP-dependent manner, the activated ubiquitin is subsequently transferred to an E2 ubiquitin-conjugating enzyme, and finally an E3 ubiquitin ligase catalyzes the transfer of ubiquitin from the E2 enzyme to a target protein (Hershko and Ciechanover, 1998). A particular E3 ligase involved in the progression of the cell cycle is the anaphase-promoting complex/cyclosome (APC/C, see (Heyman and De Veylder, 2012) for an extensive review about the plant APC/C). The APC/C is a multi-subunit complex that is strongly conserved throughout eukaryotes, with its core subunits named APC1 through APC16, of which only a few appear to be kingdom specific (Eme et al., 2011).

APC/C substrate recognition and binding is provided by two co-activators, one being the core subunit APC10 and the other being a CELL DIVISION CYCLE 20/FIZZY (CDC20/FZ) or CDC20 HOMOLOGUE 1/FIZZY-RELATED (CDH1/FZR) WD-40 domain-containing protein (Tarayre et al., 2004; da Fonseca et al., 2011; Kevei et al., 2011; Zhou et al., 2016). These activator proteins recruit the APC/C-ubiquitination targets through recognition of conserved amino acid motifs called degrons, such as the Destruction box (D-box), with the consensus sequence RxxLxxxxN, or the KEN- or GxEN-boxes (Pfleger and Kirschner, 2000; De Veylder et al., 2007; Heyman and De Veylder, 2012).

The CDC20/FZ-type activator is present in six copies in Arabidopsis, of which only CDC20.1 and CDC20.2 are thought to be functional, whereas the CDH1/FZR-type, also known as CELL CYCLE SWITCH 52 (CCS52), is present in three copies, two A-types (*CCS52A1* and *CCS52A2*) and one B-type (*CCS52B*) (Tarayre et al., 2004; Kevei et al., 2011). Both *CDC20* genes and *CCS52B* show a similar cell cycle-dependent expression, peaking from the G2- to M-phase, whereas their protein levels are temporally controlled by nuclear mRNA sequestration until breakdown of the nuclear envelope at the prometaphase, indicating that they both play a role in the progression of mitosis (Tarayre et al., 2004; Fülöp et al., 2005; Kevei et al., 2011; Yang et al., 2017). In contrast, both *CCS52A* isoforms are mainly expressed from late M-phase to G2 (Kevei et al., 2011). Plants with a diminished activity of either CCS52A isoform show a reduction in leaf cell size, accompanied by a decrease in the DNA ploidy level of these cells, while the opposite holds true for plants overexpressing either isoform (Lammens et al., 2008; Boudolf et al., 2009; Larson-Rabin et al., 2009; Baloban et al., 2013; Heyman et al., 2017). Their only partially overlapping tissue-specific expression patterns hint to both isoforms performing specific functions (Baloban et al., 2013). Correspondingly, *ccs52a1* mutants show a reduction in trichome branch number and an increased cell number within their root meristem, while *ccs52a2* mutants show unprogrammed and aberrant cell divisions in the quiescent (QC) and organizing (OC) centers of respectively the root and the shoot, accompanied by an overall dwarf growth, including a short root length and a strong reduction in the growth of inflorescences (Boudolf et al., 2009; Vanstraelen et al., 2009; Kasili et al., 2010; Liu et al., 2012; Heyman et al., 2017).

Although many plant proteins are assumed to be APC/C-ubiquitination targets, as the motifs necessary for their recognition by the APC/C are present in their amino acid sequence, only a limited set of APC/C substrates have been thoroughly characterized. Cyclins (CYCs), a family of proteins that, together with their cyclin-dependent kinase (CDK) binding partners, promote cell cycle progression through phosphorylation of target proteins, are known from research in animals and yeast to be APC/C substrates (Peters, 2002). Various A- and B-type CYCs have indeed been implicated in being targeted by the APC/C in plants, with evidence being most comprehensive for CYCA2;3 (Boudolf et al., 2009; Heyman et al., 2011; Eloy et al., 2012; Xu et al., 2016), CYCA3;4 (Fülöp et al., 2005; Willems et al., 2020), and CYCB1;1 (Kwee and Sundaresan, 2003; Fülöp et al., 2005; Rojas et al., 2009; Eloy et al., 2011; Zheng et al., 2011; Wang et al., 2012; Wang et al., 2013; Guo et al., 2016). Other cell division-related proteins characterized to be APC/C substrates include the cell wall biosynthesis gene CELLULOSE SYNTHASE LIKE-D 5 (CSLD5) (Gu et al., 2016), the rice protein ROOT ARCHITECTURE ASSOCIATED 1 (RAA1, known as FLOWERING-PROMOTING FACTOR 1 [FPF1] in Arabidopsis) that interacts with the spindle and inhibits metaphase-to-anaphase transitions (Ge et al., 2004; Han et al., 2008; Xu et al., 2010), and the Arabidopsis PATRONUS1 (PANS1) and rice RICE SALT SENSITIVE 1 (RSS1) proteins that are essential for sister chromatid cohesion during the early stages of cell division (Ogawa et al., 2011; Cromer et al., 2013; Juraniec et al., 2016; Cromer et al., 2019). Several other proteins indirectly related to the cell division process have been implicated to be APC/C targets as well, including the DSRNA-BINDING PROTEIN 4 (DRB4) involved in RNA silencing (Marrocco et al., 2012), the transcription factor ETHYLENE RESPONSE FACTOR 115 (ERF115) that controls QC divisions in the root tip (Heyman et al., 2013), the rice transcription factor MONOCULM 1 (MOC1, called SCARECROW-LIKE 18/LATERAL SUPPRESSOR [SCL18/LAS] in Arabidopsis) involved in shoot branching (Lin et al., 2012; Xu et al., 2012; Lin et al., 2020), the rice homolog of the SHORT ROOT (OsSHR) transcription factor involved in root growth (Lin et al., 2020), the rice RCAR family of abscisic acid receptors (Lin et al., 2015), and the DDR-complex subunit DEFECTIVE IN MERISTEM SILENCING 3 (DMS3) involved in the silencing of transposable elements (Zhong et al., 2019).

In this work, we exploited the outspoken short root phenotype of *CCS52A2* knock-out plants to identify novel APC/C^CCS52A2^-targets through an ethyl methanesulfonate (EMS) suppressor screen. We show that one of the identified revertants encodes a previously undescribed plant lineage-specific cell cycle protein that plays a putative role in microtubule dynamics and is targeted for destruction by APC/C^CCS52A2^.

## RESULTS

### Identification of *pkn1* as a *ccs52a2-1* Suppressor Mutant

Compared to the wild type, the *ccs52a2-1* mutant shows a strong primary root growth inhibition phenotype, especially the first days following germination, with a partial recovery during later development, reaching a total root length of 36% of that of wild-type plants at 9 days after stratification (DAS; Figures 1A, 1B, 1E and 1F, Supplemental Fig. S1A) (Vanstraelen et al., 2009; Heyman et al., 2013; Willems et al., 2020). This short root phenotype was used to identify putative targets of the APC/C^CCS52A2^ ubiquitin ligase complex through an EMS mutagenesis revertant screen. Out of a total of 260 initially identified revertants, 33 were confirmed in the next generation. Among these, one revertant mutation, named *pikmin 1-1* (*pkn1-1*), yielded a partial recovery in root length (Fig. 1C, Supplemental Fig. S1A). Compared to the *ccs52a2-1* mutant, the *pkn1-1 ccs52a2-1* double mutant showed improved root growth over almost the complete measured time course, being most outspoken at younger stages and with no significant difference at 9 DAS, reaching a total root length of 69% of that of the wild type (Fig. 1E and 1F).

**Figure 1.**
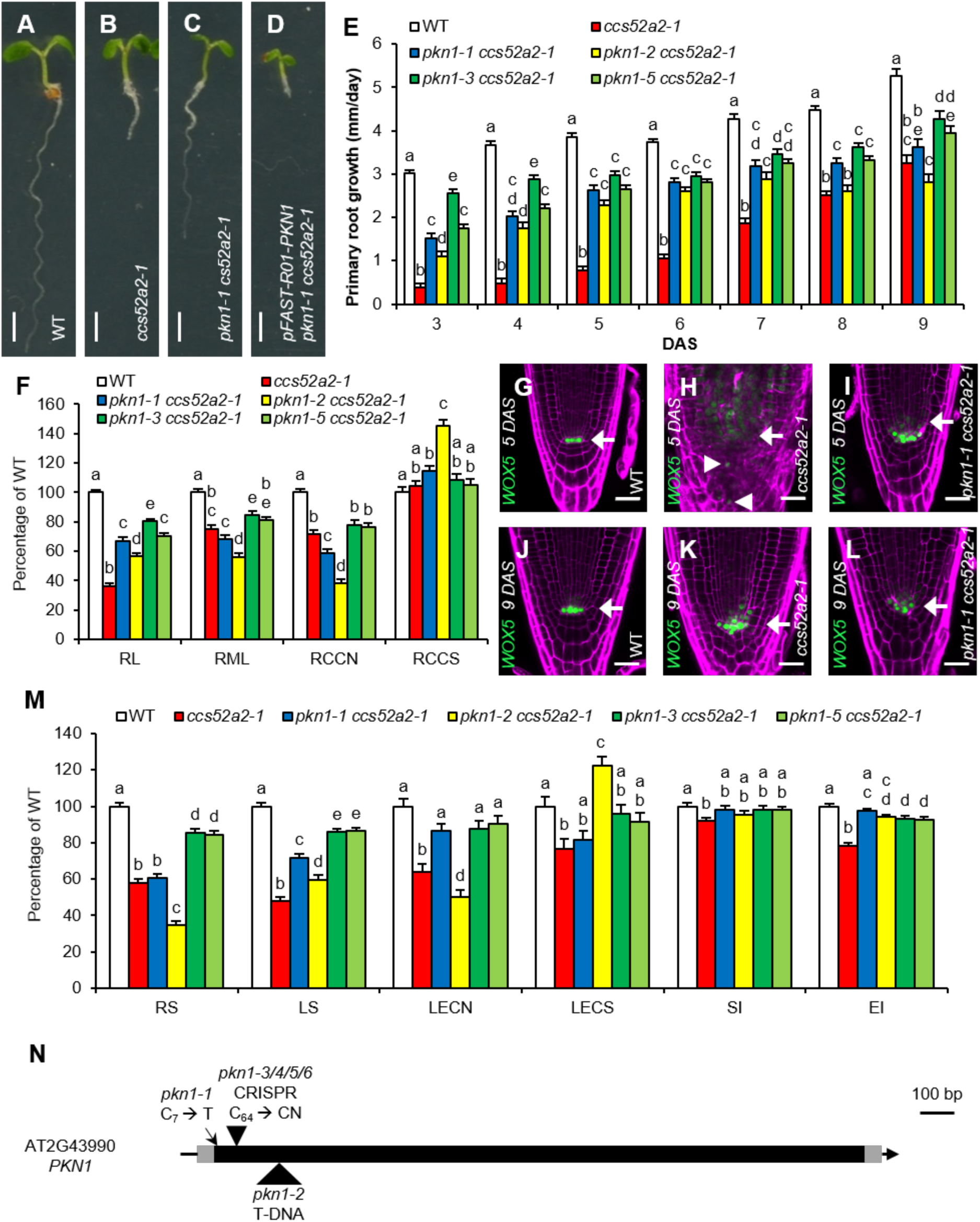
The *pkn1-1* Mutation Rescues the *ccs52a2-1* Short Root Phenotype. **(A-D)** Representative seedlings of the wild type (WT) **(A)**, *ccs52a2-1* **(B)** and *pkn1-1 ccs52a2-1* without **(C)** and with **(D)** the *pFAST-R01-PKN1* complementation construct at 5 DAS. Scale bars represent 1 mm. **(E, F and M)** Growth characteristics of WT, *ccs52a2-1* and the double mutants *pkn1-1 ccs52a2-1, pkn1-2 ccs52a2-1*, *pkn1-3 ccs52a2-1* and *pkn1-5 ccs52a2-1*. Primary root growth from 3 to 9 DAS **(E)**. Phenotypes of the primary root at 9 DAS **(F)**. RL, root length; RML, root meristem length; RCCN, root cortical cell number; RCCS, root cortical cell size. Phenotypes of the shoot and the first leaf pair at 21 DAS **(M)**. RS, rosette size; LS, leaf size; LECN, leaf epidermal cell number; LECS, leaf epidermal cell size; SI, stomatal index; EI, endoreplication index. Error bars represent standard error (n ≥ 9). Letters on the bars indicate statistically different means (P < 0.05, mixed model analysis, Tukey correction for multiple testing). **(G-L)** Representative confocal images of *proWOX5:GFP-GUS* expressing WT **(G & J)**, *ccs52a2-1* **(H & K)** and *pkn1-1 ccs52a2-1* **(I & L)** primary root tips at 5 **(G-I)** and 9 DAS **(J-L)**. The GFP signal is shown in green, while cell walls are visualized through propidium iodide staining (magenta). Arrows indicate the position of the QC, while ectopic *WOX5* expression in *ccs52a2-1* is indicated by arrowheads. Scale bars represent 25 µm. **(N)** The gene structure of the *PKN1* gene (*AT2G43990*), made up of a single exon, showing the location of the EMS mutation (arrow), T-DNA insertion (large arrowhead), and CRISPR mutations (small arrowhead). The gray and black boxes represent the untranslated regions and the coding sequence, respectively, while the lines represent intergenic sequences.

As previously described, the strongly reduced root growth in the *ccs52a2-1* mutant is accompanied by a severely disorganized root apical meristem stem cell niche (SCN) (Vanstraelen et al., 2009; Willems et al., 2020), as visualized through the use of a *proWOX5:GFP-GUS* transcriptional reporter that predominantly marks the QC stem cells. In the *ccs52a2-1* background, *WOX5* expression was detected in an expanded area of the disorganized QC and SCN during early development (at 5 DAS), as well as in differentiated columella cells and higher up in the root (Figures 1G and 1H, Supplemental Fig. S1B). At a later developmental stage (at 9 DAS), *WOX5* expression got confined to the SCN, coinciding with improved root growth and meristem organization (Figures 1J and 1K). Compared to the *ccs52a2-1* mutant, the *pkn1-1 ccs52a2-1* double mutant showed a substantially improved meristem organization at 5 DAS, accompanied by the exclusive expression of *WOX5* in the SCN, but no differences were observed at 9 DAS (Figures 1I and 1L, Supplemental Fig. S1B).

The root meristem length was measured at 9 DAS, and only reached 75% of that of the wild-type in *ccs52a2-1*, caused by a reduction in cell number, whereas cell size remained the same (Fig. 1F). In the *pkn1-1 ccs52a2-1* double mutant, meristem length and cell number did not show improvement, but were even more reduced compared to that of *ccs52a2-1*, while cell size was slightly increased (Fig. 1F), reflecting their similar root growth rates at 9 DAS (Fig. 1E).

For the shoot tissue, recovery of the *ccs52a2-1* phenotype was detected for some but not all analyzed parameters at 21 DAS (Fig. 1M). No difference in projected rosette size could be observed between *ccs52a2-1* and the double mutant, with both reaching only 58% and 61% of wild-type size, respectively (Fig. 1M). Partial recovery was observed, however, in the size of the first leaf pair, with a reduction seen in *ccs52a2-1* to 48% and the double mutant to 72% of the wild-type leaf size (Fig. 1M). The leaf growth recovery phenotype appeared to be mostly driven by a recovery in cell number, with *ccs52a2-1* showing a reduction to 64% and *pkn1-1 ccs52a2-1* to 86% of the wild-type epidermal cell number (Fig. 1M) and no significant difference in epidermal cell size, with a reduction for *ccs52a2-1* and *pkn1-1 ccs52a2-1* to 77% and 82% of the wild type, respectively (Fig. 1M). The reduction in number of stomata per pavement cell (stomatal index or SI) compared to the wild type was slightly greater in the *ccs52a2-1* mutant, with that of the double mutant being intermediate, although the differences were only minor (Fig. 1M). DNA ploidy levels, as represented by the endoreplication index (EI), were reduced in the *ccs52a2-1* mutant to 78% of the wild type. A full recovery could be observed for the double mutant, with an EI not significantly different from that of the wild type (Fig. 1M).

### Identification of the *pkn1* Mutant Gene

To identify the mutant gene underlying the *pkn1-1* mutation, a mapping scheme was set up wherein the *pkn1-1 ccs52a2-1* mutant was backcrossed with the original *ccs52a2-1* parental line and subsequently self-pollinated. In the resulting 1-in-4 segregating F2 mapping population, plants with the revertant phenotype were selected and pooled for gene mapping through next-generation sequencing, using the EMS-generated single-nucleotide polymorphisms (SNPs) as *de novo* mapping markers. Plotting the mutant allele frequency (the ratio of the number of mutant alleles over the number of wild-type alleles, also known as concordance) of the identified EMS-specific SNPs in function of their location in the genome revealed a peak at the end of chromosome 2 and subsequently an interval was selected for detailed analysis (from 17 Mbp up to the end of chromosome 2, Supplemental Fig. S2). After filtering for mutations with a concordance above 0.8 and filtering out intergenic or intronic mutations, four candidate genes were retained: *AT2G43190*, *AT2G43990*, *AT2G45360* and *AT2G46850* (Supplemental Table S1). *AT2G43190* encodes POP4, a subunit of the ribonuclease P complex involved in rRNA maturation (Gutmann et al., 2012). Because this process is deemed unlikely to play a role in *ccs52a2-1* phenotype recovery, this gene was not considered for further characterization. To identify the correct revertant gene, a SNP-genotyping PCR for the three remaining candidate mutations was performed on 133 plants with the long root phenotype from the segregating population used in the mapping of *pkn1-1 ccs52a2-1*. In this way, two plants could be isolated that were homozygous for the mutation in *AT2G43990* but contained a wild-type allele for *AT2G46850* or *AT2G45360*, indicating that the mutation in either of the latter two genes was unlikely to be responsible for the recovery phenotype. This was confirmed in the next generation, where all progeny of both plants showed the long root phenotype, and while the wild-type allele of *AT2G43990* could not be detected, the mutation in *AT2G45360* was found to be segregating in one population, while that in *AT2G46850* was segregating in both. This proved that the revertant phenotype is not correlated with the EMS mutation in either *AT2G45360* or *AT2G46850*, ruling them out and leaving *AT2G43990* as the primary candidate for the actual revertant gene. As *AT2G43990* is an unknown gene and had not been named previously, it was chosen to keep the name *PIKMIN1* (*PKN1*). The *PKN1* gene consists of a single 1899-bp exon, with the EMS mutation (C_7_ → T) changing the third codon from one encoding Arginine (CGA) to a STOP codon (TGA) (Fig. 1N, Supplemental Table S1). Furthermore, the first few subsequent ATG codons, representing alternative starting points for translation initiation, are out-of-frame, strongly indicating that the EMS mutation represents a complete knock-out of the *PKN1* gene.

Transformation of a complementation construct containing a functional copy of the *PKN1* promoter and gene (*pFAST-R01-PKN1*) into the *pkn1-1 ccs52a2-1* mutant confirmed that the *pkn1-1* mutation in *AT2G43990* was responsible for the recovery of the *ccs52a2-1* root growth phenotype, as out of the 171 transformants obtained, 168 again showed a stunted root growth phenotype (Fig. 1D). Remarkably, many transformants grew worse than *ccs52a2-1* plants, indicating that the growth phenotype of the *ccs52a2-1* plants strongly depends on PKN1 abundance and that timely breakdown of PKN1 might be essential for proper plant development.

### *PKN1* Is an Embryophyte-Specific Gene Present as a Single-Copy in Most Plant Species

A search for potential homologs did not identify any closely related gene in Arabidopsis. In fact, *PKN1* appears to be a single-copy gene in most plant species (Supplemental Data 1), as can be deduced using the PLAZA database for comparative plant genomics (https://bioinformatics.psb.ugent.be/plaza), with the *PKN1* orthologs being part of homolog group HOM04D007290 in dicotyledons (Dicots PLAZA 4.5) and HOM04x5M006718 in monocotyledons (Monocots PLAZA 4.5). BlastP searches further yielded potential *PKN1* orthologs within the lycophyte lineage, represented by *Selaginella moellendorffii*, the liverwort lineage, represented by *Marchantia polymorpha*, and the moss lineage, represented by *Physcomitrium patens* (Supplemental Data 1). No *PKN1* orthologs could be identified outside of the embryophyte clade, indicating that *PKN1* is a land plant-specific gene.

As no domain of known function is present in any PKN1 ortholog, the MEME suite (https://meme-suite.org/) was used to identify amino acid motifs conserved throughout the PKN1 ortholog lineage. Among the ten best computationally identified conserved motifs, of which motifs 1 through 8 are present in Arabidopsis PKN1 (AtPKN1), none had an annotated function (Fig. 2). Among all investigated species, only motifs 3 to 5 are conserved, with motif 5 having a minimal D-box sequence (RxxL, Supplemental Fig. S3). Within AtPKN1, a KGIL rather than the RxxL sequence is present in motif 5. Previous reports have shown that it is possible for a KxxL motif to insert in the binding pocket of APC/C (He et al., 2013; Qin et al., 2016), suggesting the KGIL in AtPKN1 motif 5 might still be recognized by the APC/C. Interestingly, all PKN1 sequences hold multiple putative APC/C recognition motifs (D-, KEN and GxEN-boxes), indirectly suggesting that PKN1 might be a bona fide APC/C target (Fig. 2). Additionally, conserved Ser-Pro (SP) or Thr-Pro (TP) can be seen in motifs 3, 6, 7, and 8 (Supplemental Fig. S3), representing potential sites for phosphorylation by CDKs or mitogen-activated protein kinases. In conclusion, the amino acid sequence analysis hints at PKN1 being a conserved protein that is potentially regulated in a cell cycle-dependent manner through ubiquitination by the APC/C and phosphorylation.

**Figure 2:**
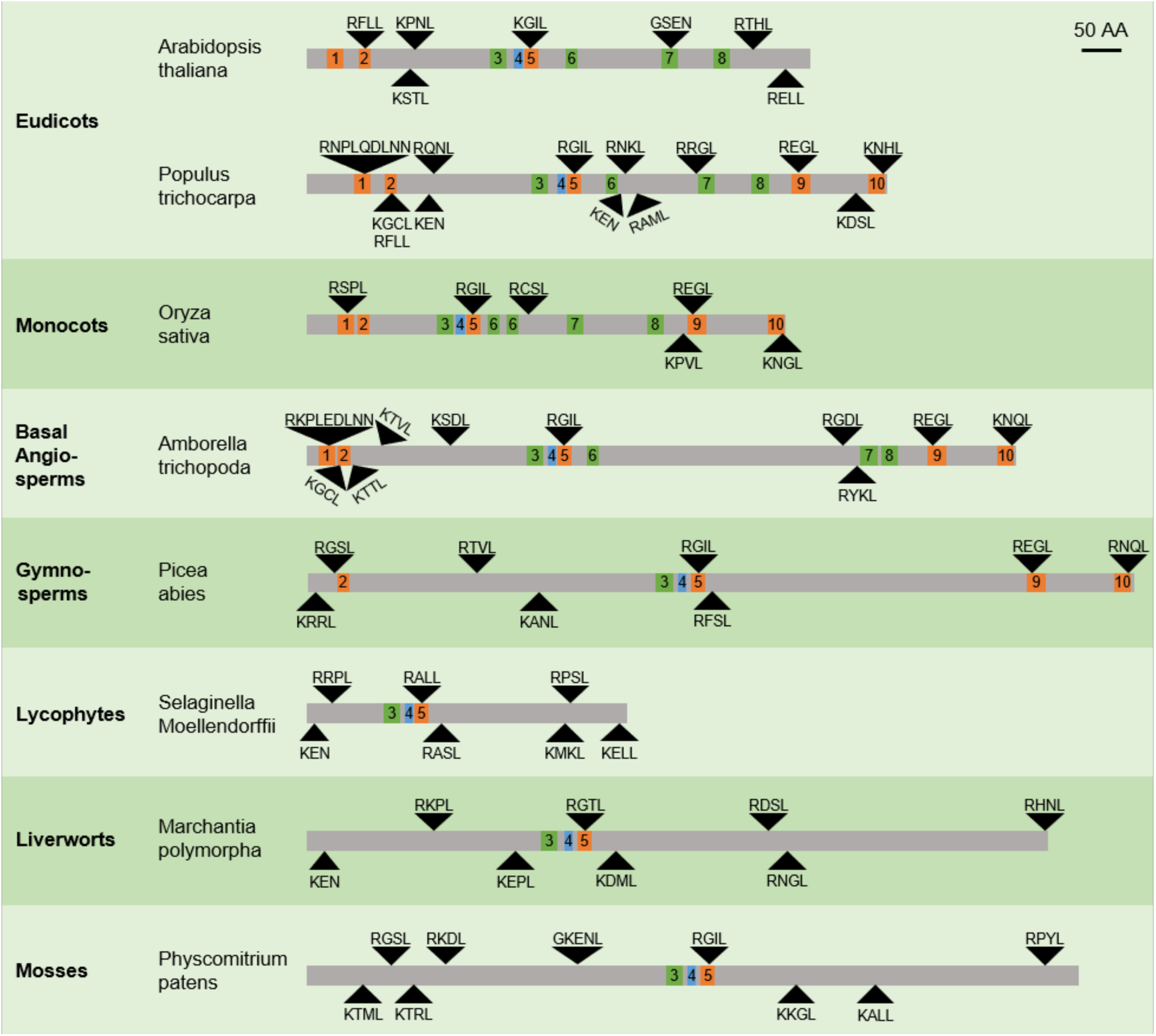
Conserved Motifs in the PKN1 Protein. The Arabidopsis thaliana PKN1 protein and its orthologs throughout the plant kingdom, showing the location of the different conserved motifs identified (colored boxes) and of the D- (showing both RxxL and KxxL sequences), KEN- or GxEN-boxes (black triangles). Motifs often correlating with a D-box are shown in orange, those correlating with a possible phosphorylation site in green, and those without clear function in blue.

### PKN1 Is Rate-Limiting for Cell Division in the Root Meristem

To determine the effects of the lack of PKN1 on plant development, the *ccs52a2-1* mutation was crossed out of the *pkn1-1 ccs52a2-1* double mutant using Col-0, creating the *pkn1-1* single mutant. Although root length of *pkn1-1* did not differ from that of the wild type at 9 DAS, the root meristem length was reduced by 32% and the root meristem cortical cell number was reduced by 39%, compensated by a small 13% increase in root cortical cell size (Fig. 3A). On the level of the shoot, no significant differences could be observed for any of the investigated characteristics, except for a minor 10% increase in mature leaf size (Fig. 3A).

**Figure 3:**
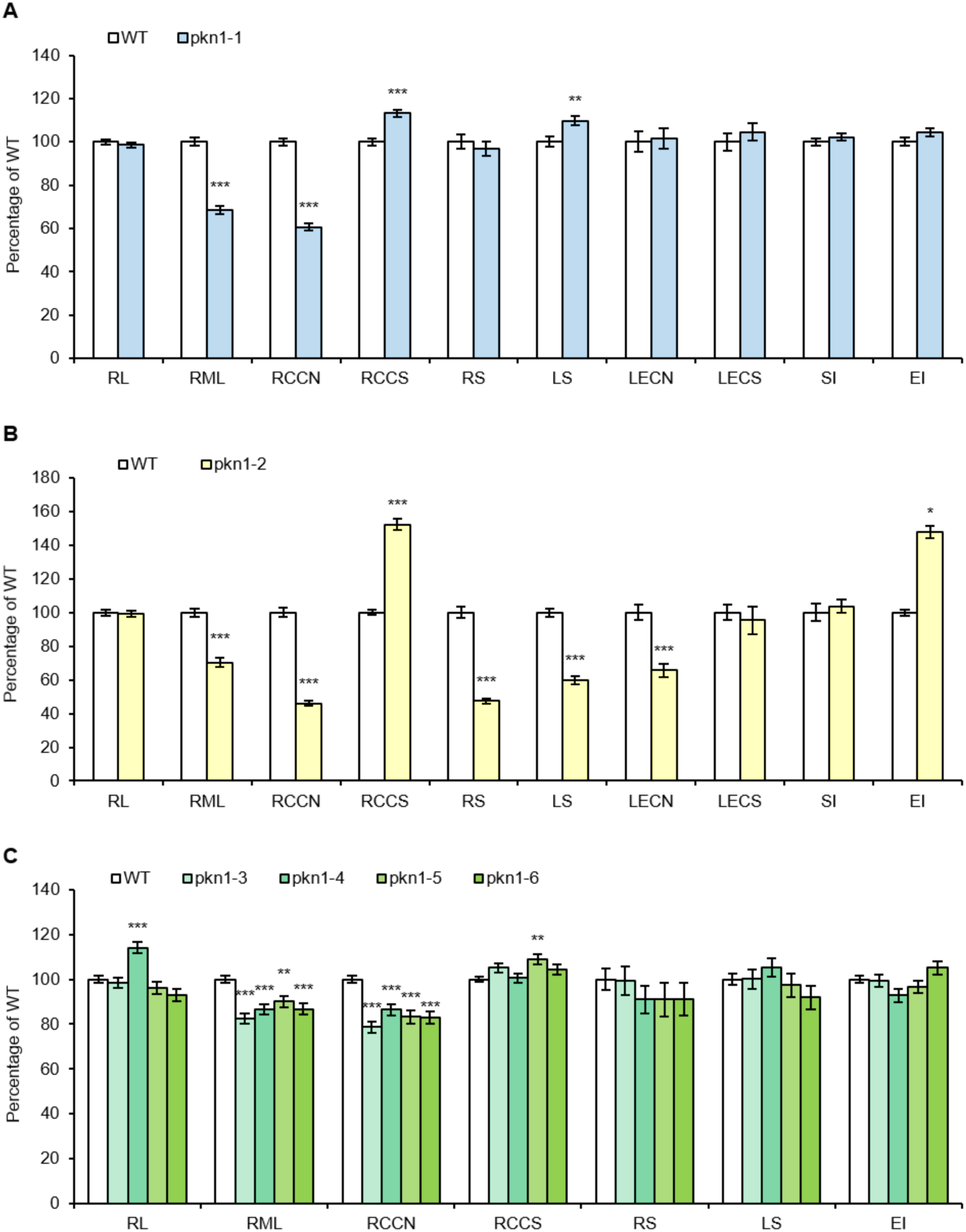
Characteristics of *pkn1* Knock-Out Mutants. **(A)** Phenotypes of the *pkn1-1* mutant in the root at 9 DAS and the shoot at 21 DAS; RL: root length (n > 34), RML: root meristem length (n > 29), RCCN: root cortical cell number (n > 29), RCCS: root cortical cell size (n > 29), RS: rosette size (n > 38), LS: leaf size (n = 38), LECN: leaf epidermal cell number (n = 10), LECS: leaf epidermal cell size (n =10), SI: stomatal index (n = 10), EI: endoreplication index (n = 10). Error bars represent standard error and stars represent means significantly different from the wild-type (WT) mean (Mixed model analysis, ** p < 0.01, *** p < 0.001). **(B)** Phenotypes of the *pkn1-2* T-DNA mutant in the root at 9 DAS and the shoot at 21 DAS; RL (n > 22), RML (n = 24), RCCN (n = 24), RCCS (n = 24), RS (n > 28), LS: leaf size (n > 14), LECN: leaf epidermal cell number (n = 3), LECS: leaf epidermal cell size (n = 3), SI: stomatal index (n = 3), EI: endoreplication index (n = 2). Error bars represent standard error and stars represent means significantly different from the WT mean (Two-sample *t*-test, * p < 0.05, *** p < 0.001). **(C)** Phenotypes of the *pkn1-3, pkn1-4, pkn1-5* and *pkn1-6* CRISPR mutants in the root at 9 DAS and the shoot at 21 DAS; RL (n > 24), RML (n > 17), RCCN (n > 17), RCCS (n > 17), RS (n > 11), LS (n = 11), EI (n = 6). Error bars represent standard error and stars represent means significantly different from the WT mean (Mixed model analysis with Dunnett correction for multiple testing, ** p < 0.01, *** p < 0.001).

To confirm the *pkn1-1* phenotypes, an independent mutant was isolated from the SALK T-DNA insertion collection (SALK_047285) and named *pkn1-2*. Knock-out was confirmed using RT-PCR (Supplemental Fig. S4) and the T-DNA insertion location was determined through Sanger sequencing to be in the coding sequence of *PKN1* behind base C^189^ (Fig. 1N). Identical to *pkn1-1*, the root length of *pkn1-2* did not differ from the wild type, and the root meristem length was reduced by 30% (Fig. 3B). However, both the decrease in cortical cell number (minus 54% compared to the wild type) and the increase in cell size (plus 52% compared to the wild type) were more drastic in *pkn1-2* than in *pkn1-1*. Contrastingly to *pkn1-1*, the *pkn1-2* mutant did show phenotypes in the shoot. At 21 DAS, the rosette size of *pkn1-2* plants was reduced by 53%, reflected by a reduction in leaf size of 40% in the first leaf pair (Fig. 3B). This reduction was due to a 34% decrease in epidermal cell number, with cell size and SI remaining unchanged (Fig. 3B). The level of endoreplication, reflected by the EI, was also increased by 48% compared to the wild type (Fig. 3B). The reduction in cell number and increase in endoreplication indicate that cells exited the proliferation phase and entered differentiation prematurely in *pkn1-2* mutant plants.

To confirm *PKN1* as the correct revertant gene, the *pkn1-2* mutant was introgressed into the *ccs52a2-1* background, resulting in the *pkn1-2 ccs52a2-1* double mutant. Root growth of the *pkn1-2 ccs52a2-1* double mutant showed partial recovery of *ccs52a2-1* root growth, especially early during development, but was less than that of *pkn1-1 ccs52a2-1* (Fig. 1E). Concurrently, the root meristem at 9 DAS of the double mutant was shorter than that of both *ccs52a2-1* and *pkn1-1 ccs52a2-1*, due to a further reduction in cell number accompanied by a greater increase in cell size (Fig. 1F). Shoot phenotypes of *pkn1-2 ccs52a2-1* were reminiscent of those of the *pkn1-2* single mutant, showing a strongly reduced rosette size being even smaller than that of *ccs52a2-1*, while leaf size was only slightly increased compared to that of *ccs52a2-1* (Fig. 1M). Leaf epidermal cell number was also reduced compared to that of *ccs52a2-1*, but cell size was increased compared to the wild type, explaining the partial leaf size recovery (Fig. 1M). Additionally, the SI and EI of *pkn1-2 ccs52a2-1* was identical to that of *pkn1-1 ccs52a2-1* (Fig. 1M).

To address the discrepancy between the *pkn1-1* and *pkn1-2* phenotypes, independent *PKN1* mutant lines were generated using the CRISPR/Cas9 technology. Two different guide RNAs were cloned into the same vector, with the expected cut sites in the beginning of the gene, after respectively base pair 64 and 286 of the PKN1 CDS. For each of six different T1 primary transformants, seven to eight plants without the CRISPR/Cas9 construct were selected in the segregating T2 generation and screened for mutations at the two cut sites using SNP genotyping. Subsequently, the presence and the exact nature of the mutations were determined by Sanger sequencing. This showed that editing at the first cut site was highly efficient, with many single base pair INDELs found, but no mutations were identified at the second cut site. Some of the screened plants were already homozygous, and of those, four were retained, being *pkn1-3* (insertion of T), *pkn1-4* (insertion of T), *pkn1-5* (insertion of C) and *pkn1-6* (insertion of A). In each case, the insertion behind C^64^ resulted in a frame shift and premature stop after six codons, most likely leading to a complete abolishment of PKN1 function (Fig. 1N).

Identical to the *pkn1-1* mutant line, root length at 9 DAS was not significantly different for all of the CRISPR lines but *pkn1-4*, while root meristem length was reduced in all lines by 10-18% compared to the wild type, caused by a reduction in cell number by 14-21% (Fig. 3C). However, a large cell size increase, reminiscent to that of *pkn1-1* or *pkn1-2* could not be seen (Fig. 3C). Like the *pkn1-1* mutant, no difference compared to the wild type was observed for the *PKN1* CRISPR mutants regarding any of the leaf parameters tested at 21 DAS, including rosette size, size of the first leaf pair and ploidy levels (Fig. 3C). Two of the CRISPR mutants, *pkn1-3* and *pkn1-5*, were introgressed into the *ccs52a2-1* background, and both double mutants showed an equal or better recovery of *ccs52a2-1* phenotypes compared to *pkn1-1 ccs52a2-1*, confirming *PKN1* again as the true revertant gene (Figures 1E, 1F and 1M). Taken together, it seems that the *PKN1* gene plays an important role in cell proliferation in the root meristem, but not during leaf development, although its absence does partially rescue the reduced leaf cell phenotype of *ccs52a2-1*.

### Ectopic *PKN1* Expression Promotes Cell Division in the Leaf

To further investigate the potential role of PKN1 in cell division, *PKN1* overexpression lines (*PKN1^OE^*) were generated by expressing the *PKN1*-coding sequence using the strong *Cauliflower Mosaic Virus 35S* (*CaMV 35S*) promoter. Overexpression levels of *PKN1* were measured in young rosettes (10 DAS) of homozygous T3 lines and ranged between 4 and 90 times that of the wild type (Fig. 4A). For five of those lines *PKN1* expression levels were also measured in the root tip and were found to range between 5 and 16 times that of wild type (Fig. 4B). For phenotypical analysis, three lines were chosen with low (line 10.5), medium (line 7.6) and high (line 12.4) *PKN1* overexpression levels. In the shoot, all three lines showed an increased rosette size, with that of the high overexpressing line being 19% larger than that of the wild type (Fig. 4C). This was reflected in the first leaf pair, with that of the medium line being 12% and that of the high line 17% larger than that of the wild type (Fig. 4C). This increase in leaf size was caused by an increase in cell number, with the leaf epidermal cell number being increased between 10 and 17% for the three lines, while no difference was found for epidermal cell size (Fig. 4C). Interestingly, this increase in cell number was mostly restricted to the stomata for the low and medium line, while the high line showed a similar increase in both stomata and pavement cells, leading to a higher SI for the low and medium lines, but not for the high line (Fig. 4C). Additionally, the EI was not different compared to the wild type (Fig. 4C). In the root, *PKN1* overexpression did not affect growth, although a small, but significant, increase in root length was seen for the low and high line, accompanied by a small, but not statistically significant, increase in root meristem length and cell number (Fig. 4D).

**Figure 4:**
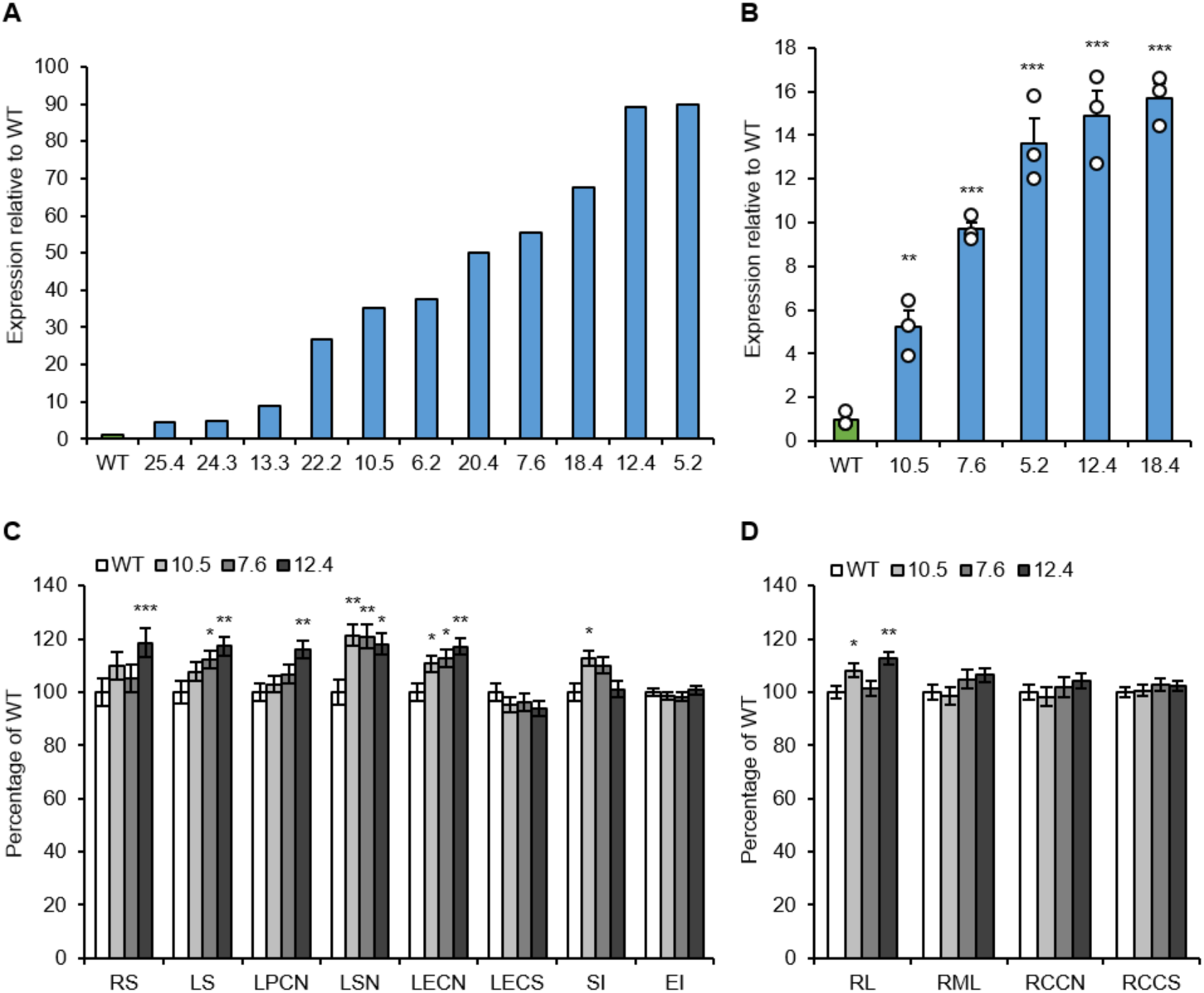
Overexpression of *PKN1.* **(A-B)** *PKN1* expression levels in *PKN1^OE^* homozygous lines of whole rosettes at 10 DAS **(A)** and of root tips at 9 DAS **(B)**. Dots represent each sample (n=3) and the bar represents the mean. Error bars represent standard error and stars represent means significantly different from the wild-type (WT) mean (Two-sample *t*-test, * p < 0.05, ** p < 0.01, *** p < 0.001). **(C-D)** Characteristics of the *PKN1^OE^* lines in the shoot at 21 DAS **(C)** and the root at 9 DAS **(D)**. RS: Rosette size (n > 11), LS: leaf size (n > 10), LPCN: leaf pavement cell number (n = 10), LSN: leaf stomatal number (n = 10), LECN: leaf epidermal cell number (n = 10), LECS: leaf epidermal cell size (n =10), SI: stomatal index (n = 10), EI: endoreplication index (n = 5), RL: root length (n > 13), RML: root meristem length (n > 10), RCCN: root cortical cell number (n > 10), RCCS: root cortical cell size (n > 10). Error bars represent standard error and stars represent means significantly different from the WT mean (ANOVA with Dunnett correction for multiple testing, * p < 0.05, ** p < 0.01, *** p < 0.001).

In conclusion, analysis of *PKN1* overexpression lines suggests a rate-limiting role for PKN1 in promoting cell division. The fact that the impact of increased PKN1 levels on cell number is mostly observed in the shoot might be due to an already higher level of PKN1 in the root tip than in the shoot in the wild-type plants (Supplemental Fig. S5).

### *PKN1* Localizes at Microtubular Structures

To further investigate the possible function of the PKN1 protein, a GUS translational reporter construct was made, *proPKN1:PKN1-GUS*. The PKN1 protein appeared to accumulate in dividing tissues throughout the plant, including the shoot and root apical meristems, the lateral and adventitious root initiation sites, the developing young leaves, the stomata, the trichome socket cells, the ovary and the developing embryo (Figures 5A to 5H). These tissue-specific accumulation patterns corroborate the idea that PKN1 plays a role in cell division.

**Figure 5:**
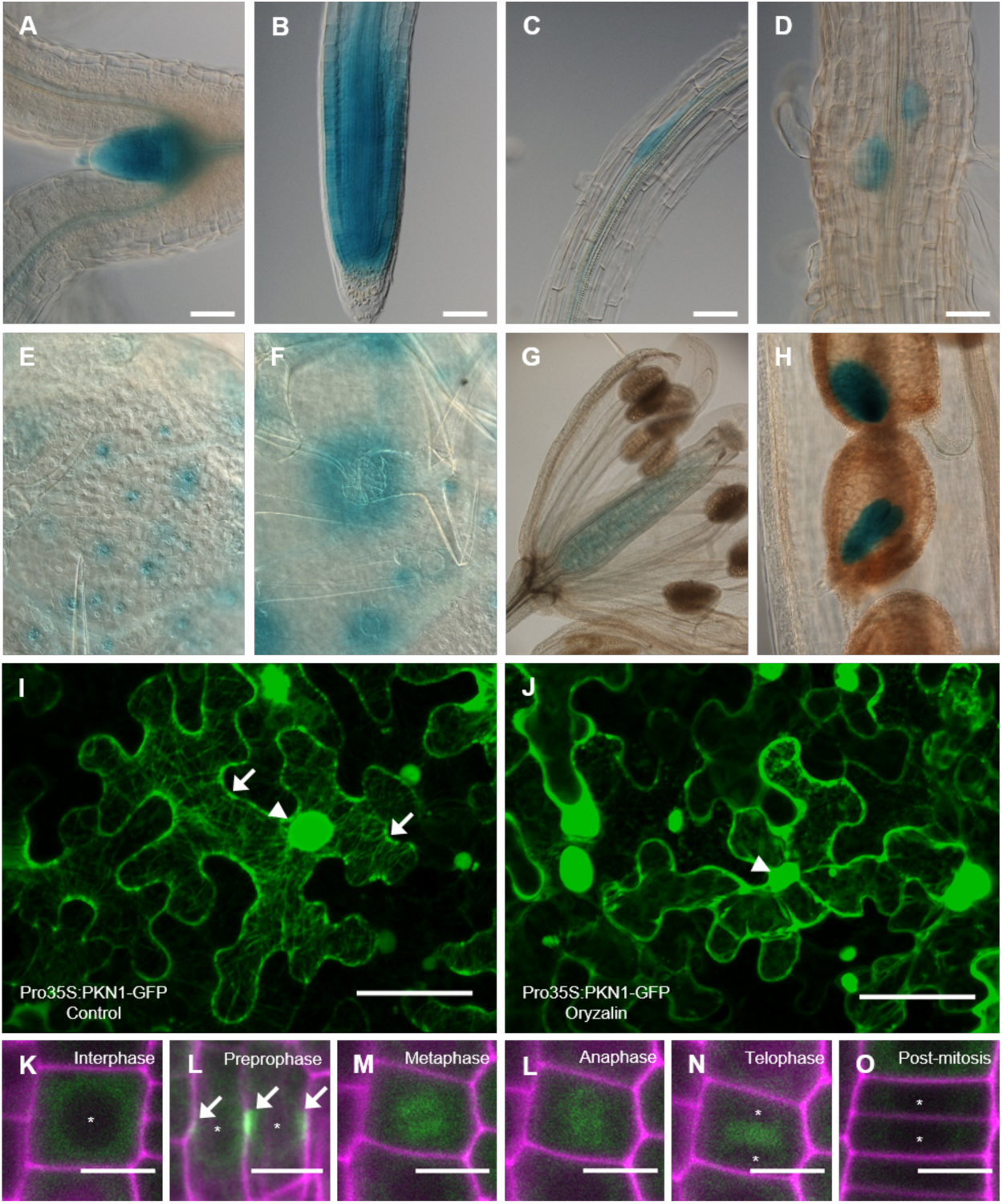
Tissue-Specific Expression Pattern and Intracellular Localization of PKN1. **(A-H)** Expression of the *proPKN1:PKN1-GUS* construct was detected in the shoot apical meristem and developing young leaves **(A)**, the root apical meristem **(B)**, the adventitious **(C)** and lateral root initiation sites **(D)**, the stomata **(E)**, the trichome socket cells **(F)**, the developing ovules **(G)** and the growing embryo **(H)**. Scale bars represent 50 µm **(A-F)**, 500 µm **(G)** or 100 µm **(H)**. **(I-J)** Transient expression of *pro35S:PKN-GFP* in tobacco leaf epidermal cells after agroinfiltration, under control conditions **(I)** or after a 1,5 h treatment with oryzalin **(J)**. In the control conditions **(I)**, the GFP signal can be seen in the cytosol at the cell membrane, at the nucleus (arrowhead) and on strands of microtubules enveloping the cell (arrows), whereas after treatment with the microtubule-destabilizing drug **(J)** the microtubule GFP localization signal disappears. Pictures are 3D projections of a Z-stack. Scale bars represent 50 µm. **(K-O)** Expression of *proPKN1:PKN1-GFP* in root meristem cells. The PKN1-GFP protein shows a cell cycle-dependent localization: during interphase **(K)** it is present in the cytosol but excluded from the nucleus (star); during preprophase **(L)** it also locates to the preprophase band (arrows); during mitosis **(M-N)** it appears to locate with the spindle microtubules **(M and L)** and the phragmoplast (**N**, stars show the newly forming nuclei); post-mitosis its signal is drastically reduced (**K**, stars show the newly formed nuclei). The GFP signal is shown in green, while cell walls are visualized through propidium iodide staining (magenta). Scale bars represent 10 µm.

To investigate the intracellular PKN1 localization pattern, GFP translational reporter constructs were generated. One construct, *pro35S:PKN1-GFP*, expressing PKN1 under the control of the *CaMV 35S* promoter, was transiently transformed into tobacco (*Nicotiana benthamiana*) leaves through agroinfiltration, while two others, *proPKN1:PKN1-GFP* and *pro35S:GFP-PKN1,* expressing the PKN1-GFP fusion under the control of its native promoter or the *CaMV 35S* promoter, respectively, were stably transformed into *Arabidopsis thaliana*. In the tobacco leaf epidermal cells, GFP signal was detected in the cytosol, both at the cell membrane and around the nucleus, as well as on cortical microtubules (Fig. 5I). Microtubule-specific localization was confirmed by the disappearance of the signal following treatment with the microtubule-depolymerizing drug oryzalin (Fig. 5J). Confocal imaging of root tips of Arabidopsis plants expressing the *proPKN1:PKN1-GFP* or *pro35S:GFP-PKN1* construct revealed that PKN1 appeared to show a cell cycle phase-dependent intracellular localization pattern (Figures 5K to 5O, Supplemental Fig. S6). In cells going through interphase, the GFP signal appeared to be confined to the cytosol (Fig. 5K, Supplemental Fig. S6). Subsequently, before mitosis, a stronger GFP signal was seen at the position of the preprophase band (Fig. 5L), while during mitosis, GFP localized at the spindle microtubuli and the expanding phragmoplast (Figures 5M to 5N, Supplemental Fig. S6). Following division, the GFP signal was strongly reduced, indicating that PKN1 might be actively broken down after completion of mitosis (Fig. 5O, Supplemental Fig. S6).

### CCS52A2 Targets PKN1 for Destruction

To investigate if PKN1 might be targeted for proteasomal destruction, the *proPKN1:PKN1-GUS* reporter line was treated with the proteasome blocker MG-132. To assure that a difference in staining intensity could be observed, GUS staining was purposefully kept short. Under control conditions, no staining could be observed in a weak *proPKN1:PKN1-GUS* line (Fig. 6A), while after an overnight treatment with MG-132, the GUS signal was visible throughout the root meristem (Fig. 6B). Similarly, a stronger *proPKN1:PKN1-GUS* line showed less signal under control conditions than following MG-132 treatment (Figures 6C and 6D). To rule out that the observed effect was due to differences on the transcriptional level, the translational reporter *pro35S:GFP-PKN1* was also tested. As with the GUS reporter, only a faint GFP signal could be observed under control conditions (Fig. 6E), whereas following MG-132 treatment, the GFP signal was intense and clearly visible (Fig. 6F).

**Figure 6:**
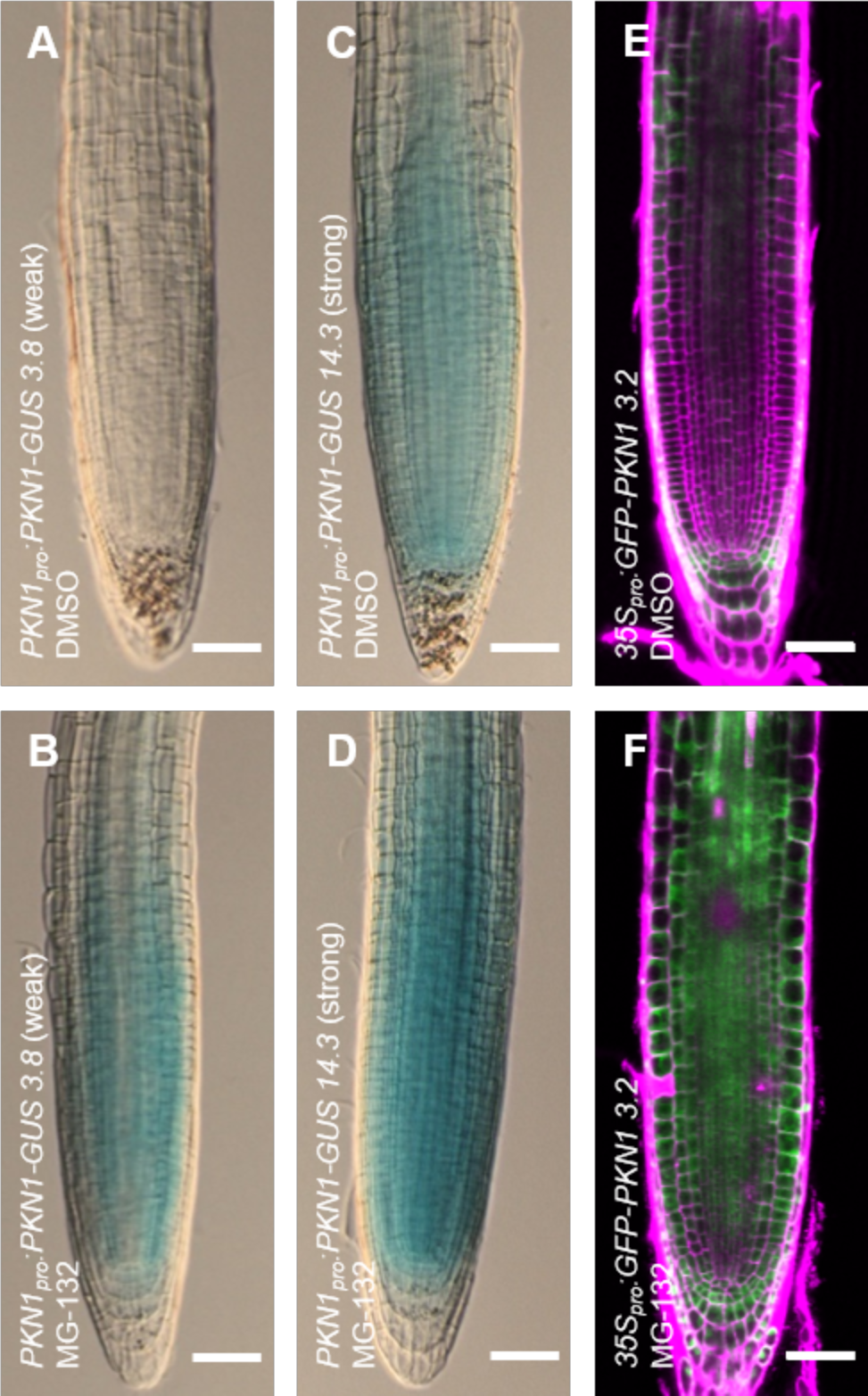
PKN1 is under control of the 26S proteasome. **(A-D)** Seedlings at 5 DAS of a weak **(A and B)** and a strong **(C and D)** *proPKN1:PKN1-GUS* line were treated overnight with DMSO control **(A and C)** or 100 µM MG-132 dissolved in DMSO **(B and D)**, and subsequently briefly stained with X-Gluc for 60 min at 37°C. Scale bars represent 50 µm. **(E-F)** Seedlings at 5 DAS of a *pro35S:GFP-PKN1* line were treated overnight with DMSO control **(E)** or 100 µM MG-132 dissolved in DMSO **(F)**, and subsequently imaged trough confocal microscopy using identical settings. GFP signal is shown in green, while cell walls are visualized through propidium iodide staining (magenta). Scale bars represent 50 µm.

To further test whether PKN1 is marked for breakdown by CCS52A2-mediated APC/C activity, the *proPKN1:PKN1-GUS* reporter line was introgressed into the *ccs52a2-1* mutant background. Interestingly, the root growth of plants homozygous for *ccs52a2-1* and containing the *PKN1-GUS* construct was dramatically reduced compared to *ccs52a2-1* plants without the construct and was often completely stalled (Figures 7A to 7E). This phenotype was probably caused by increased PKN1 protein levels, introduced by the reporter construct, again highlighting the importance in controlling PKN1 protein abundance through breakdown via CCS52A2. When comparing the PKN1-GUS activity in wild-type versus *ccs52a2-1* plants, the staining in the root meristem appeared more intense in the *ccs52a2-1* background compared to the wild-type background (Figures 7F to 7G and 7I to 7J), suggesting that PKN1 is indeed targeted for destruction by APC/C^CCS52A2^. Furthermore, meristem organization appeared even more disturbed compared to *ccs52a2-1*, and sporadically meristems were forked (Fig. 7H) or completely consumed (Fig. 7K). Taken together, these data indicate that it is essential for root meristem development that PKN1 protein abundance is controlled through destruction by APC/C^CCS52A2^.

**Figure 7:**
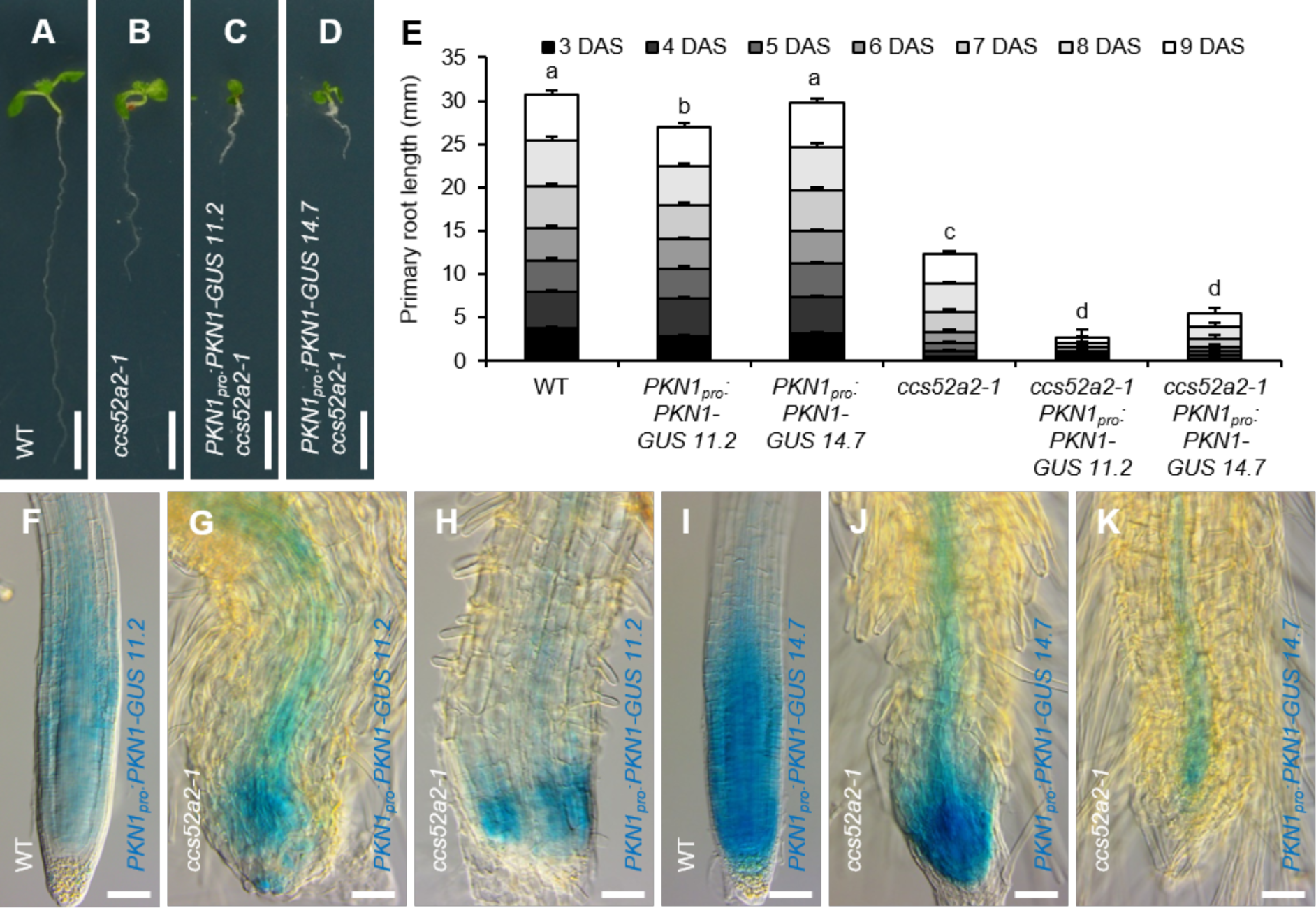
The *proPKN1:PKN1-GUS* Reporter in the *ccs52a2-1* Background. **(A-D)** Representative seedlings at 9 DAS of the wild type (WT) **(A)**, *ccs52a2-1* **(B)** and *proPKN1:PKN1-GUS* line 11.2 **(C)** or line 14.7 **(D)** in the *ccs52a2-1* background. Scale bars represent 5 mm. **(E)** Primary root length of the respective *proPKN1:PKN1-GUS* lines in the WT and *ccs52a2-1* background from 3 to 9 DAS. Bars represent estimated marginal means and bar heights were subdivided according to the measured daily growth. Error bars represent standard error (n ≥ 13), and letters indicate statistically different means for each genotype, as calculated for the total root length at 9 DAS (P < 0.05, ANOVA mixed model analysis, Tukey correction for multiple testing). **(F-K)** Histochemical GUS staining at 5 DAS of WT **(F and I)** and *ccs52a2-1* KO **(G, H, J and K)** root tips with either *proPKN1:PKN1-GUS* 11.2 **(F-H)**, or *proPKN1:PKN1-GUS* 14.7 **(I-K)** constructs in their background. Roots were stained for 1 h. Scale bars represent 50 µm.

### The PKN1 Role in Cell Division Is Conserved in *Marchantia polymorpha*

The *Marchantia polymorpha* genome holds one putative *PKN1* homologous gene (*Mp7g06500*, Supplemental Data Set 1), nominated *MpPKN1*. As observed for its Arabidopsis counterpart, *MpPKN1* expression correlated with cell division activity, as transcript levels were low in the non-dividing gemmae but strongly increased in 3-day-old dividing young gemmalings (Fig. 8A). To confirm its role during cell division, three independent CRISPR/Cas9-based knock-out lines were generated (Fig. 8B). One line (#28) holds a single base pair insertion at the first target site (after base pair G^80^), whereas the other two lines (#5 and #29) hold an inversion of different lengths of the sequence between the two target sites (in between base pairs G^81^ and C^360^). Wild-type and mutant non-dividing gemmae were indistinguishable, showing an identical epidermal cell number, epidermal cell size and total plant size (day 0 in Figures 8C to 8E). However, three days after cell cycle activation, the mutant gemmalings showed a reduced epidermal cell number compared to wild-type gemmalings (Fig. 8C), suggesting an inhibition of cell cycle activity. This was correlated with a compensatory increase in cell size, most outspoken at the apical notches (Figures 8D, 8F and 8G), leading to the total size of the mutant gemmalings remaining identical to that of the control (Fig. 8E). Likewise, the area of 28-day-old mature thalli did not differ between the different genotypes (Figures 8H to 8L).

**Figure 8.**
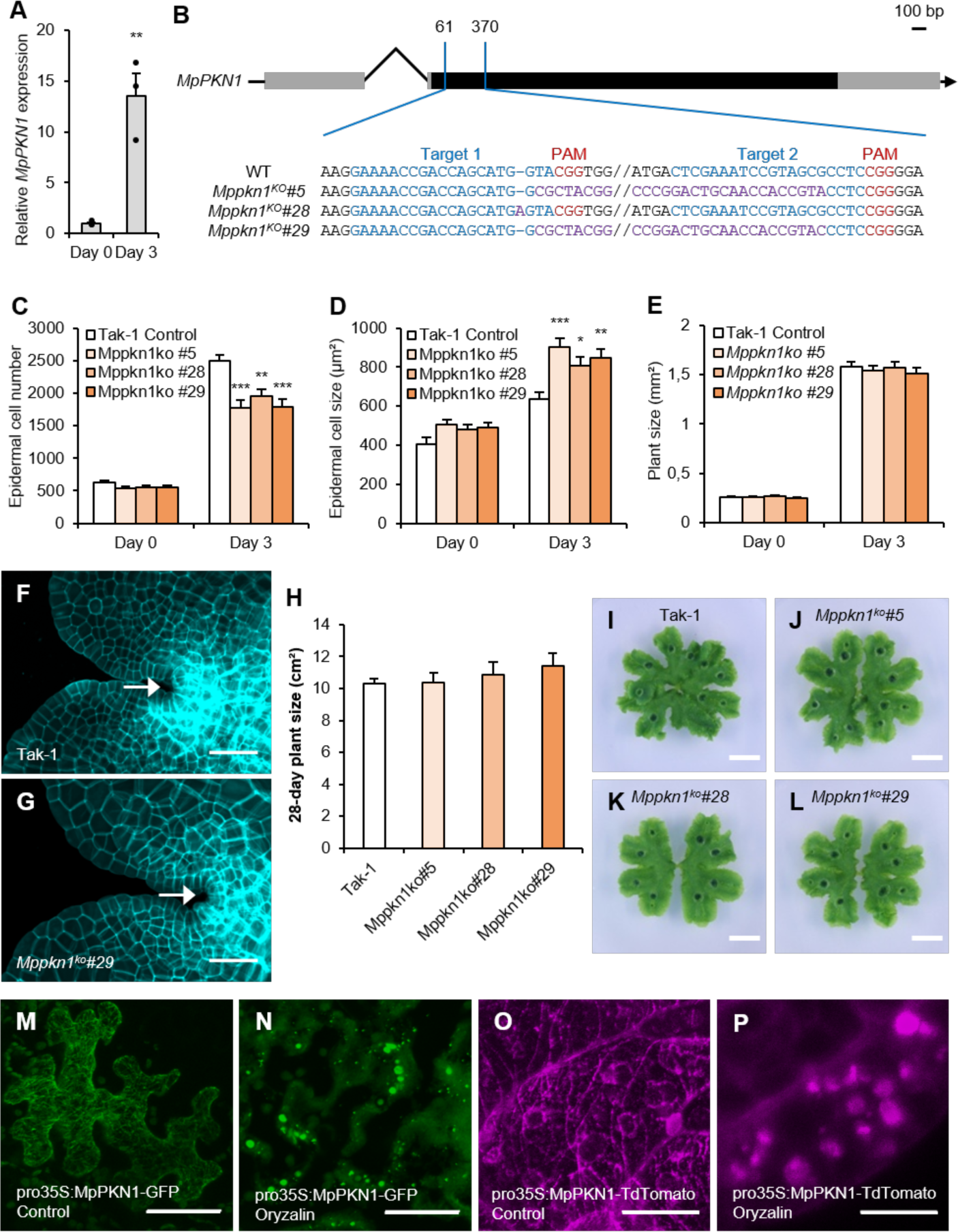
*MpPKN1* Controls Cell Division in *Marchantia polymorpha*. **(A)** Time-course of *MpPKN1* expression levels in Tak-1 gemmae (day 0) and 3-day-old gemmalings. Dots represent individual data points (n = 3). Error bars represent standard error and stars represent means significantly different from the wild-type (WT) mean (T-test with Bonferroni correction for multiple testing, ** p < 0.01). **(B)** Gene structure of the *MpPKN1* gene (Mp7g06500, Mapoly0057s0017), showing the three independent CRISPR mutations lines (*#5*, *#28* and *#29*). The gray and black boxes represent the untranslated regions and the coding sequences, respectively, while blue, red and purple sequence represents the gRNA, PAM site and inverted sequence, respectively. **(C-E)** Epidermal cell number **(C)**, epidermal cell size **(D)**, and plant size **(E)** of gemmae (day 0) and 3-day-old gemmalings of Tak-1 and *Mppkn1^ko^#5*, *#28* and *#29*. Error bars represent standard error (n ≥ 6) and stars represent means significantly different from the WT mean (ANOVA with Dunnett correction for multiple testing, * p < 0.05, ** p < 0.01, *** p < 0.001). **(F-G)** Representative images of apical notches of 3-day-old Tak-1 **(F)** and *Mppkn1^ko^#29* **(G)** gemmalings. Cell walls were stained with calcofluor white (CFW). The arrows indicate apical notches. Scale bars represent 50 µm. **(H)** Plant size of 28-day-old Tak-1 and *Mppkn1^ko^#5*, *#28* and *#29* thalli. Error bars represent standard error (n ≥ 6), no significant difference was found (ANOVA with Dunnett correction for multiple testing). **(I-L)** Aerial view of representative 28-day-old mature Tak-1 and *Mppkn1^ko^* plants. Scale bars represent 1 cm. **(M-O)** Representative confocal images of the *pro35S:MpPKN1-GFP* construct infiltrated in *N. benthamiana* **(M and N)** and the *pro35S:MpPKN1-TdTomato* construct transformed in *M. polymorpha* **(O and P)**, treated for 1 h with control **(M and O)** or 10 µM of the microtubule-depolymerizing drug oryzalin **(N and P)**. The MpPKN1 localization to the cable-like microtubules seen under control conditions is abolished after treatment with oryzalin. Scale bars represent 50 µm **(M and N)** or 20 µm **(O and P)**.

To study the intracellular MpPKN1 localization pattern, a C-terminal GFP translational reporter construct, *pro35S:MpPKN1-GFP,* was transiently transformed into tobacco (*N. benthamiana*) leaves through agroinfiltration, while a C-terminal TdTomato translational reporter construct, *pro35S:MpPKN1-TdTomato*, was stably transformed into Marchantia. For both constructs, imaging revealed fluorescent reporter signals localizing at the cortical microtubules, which could be disrupted through treatment with the microtubule-depolymerizing drug oryzalin (Figures 8M to 8P).

## DISCUSSION

### The Mutation in *PKN1* Is Responsible for the Recovery of *ccs52a2-1* Phenotypes

In this study, the previously described short root phenotype of plants lacking the APC/C-activator CCS52A2 (Vanstraelen et al., 2009; Liu et al., 2012; Baloban et al., 2013; Willems et al., 2020) was used as the basis of an EMS mutagenesis suppressor screen, aiming at the identification of potential novel APC/C-ubiquitination targets. Three lines of evidence proved that the nonsense mutation in a previously undescribed gene, which we dubbed *PKN1*, was causative for the partial recovery of the *ccs52a2-1* growth phenotypes seen in *pkn1-1 ccs52a2-1* plants. Firstly, a SNP-genotyping PCR was used to determine that two other putatively linked candidate mutations were not linked to the *ccs52a2-1* recovery phenotype, with only the mutation in *PKN1* remaining as a valid candidate. Secondly, *pkn1-1 ccs52a2-1* plants holding a *PKN1* complementation construct showed a growth identical to or worse than that of *ccs52a2-1*, not only proving that *PKN1* is indeed the correct revertant gene, but also indicating that it is essential for plant growth that PKN1 proteins are broken down in a timely manner. Finally, independent *PKN1* mutants were shown to be able to partially rescue at least some of the *ccs52a2-1* dwarf growth phenotypes, separately confirming that *PKN1* represents the true revertant gene.

### The Different *PKN1* Knock-Out Lines Show Individual and Common Phenotypes

All types of *pkn1* mutations analyzed likely completely abolish PKN1 function, due to their early positions in the coding sequence, resulting in out-of-frame translation (Fig. 1N). Despite some variation, a phenotype in common between all mutants included a shorter root meristem that could be attributed to a reduction in cell number. In the EMS-induced (*pkn1-1*) and T-DNA (*pkn1-2*) mutants, more severe phenotypes were seen compared to the CRISPR-generated mutants (*pkn1-3/4/5/6*). This might be explained by the presence of additional background EMS mutations or the T-DNA insertion affecting a nearby gene (*AT2G43980*, encoding the enzyme inositol 1,3,4-trisphosphate 5/6-kinase 4), respectively. In contrast, the highly specific nature of the CRISPR gene editing technology, resulting here in a single base pair insertion, makes the presence of other mutations less likely, thus making the CRISPR mutants the most suited for future analysis of *PKN1* function. Similarly, the phenotypical differences between the single *PKN1* mutants are reflected in the *pkn1 ccs52a2-1* double mutants as well, with the recovery of *ccs52a2-1* phenotypes being more outspoken in the CRISPR double mutants, again indicating that growth in the *pkn1-1* and *pkn1-2* lines is additionally influenced by other factors besides the lack in PKN1 function. Taken together, all *pkn1 ccs52a2-1* double mutants show a better root growth compared to that of *ccs52a2-1*, especially in the first days post-germination, which can be ascribed to a greatly improved root SCN organization. At later growth stages, the difference in root growth between the *cc52a2-1* single and *pkn1 ccs52a2-1* double mutants becomes less outspoken, which can be explained by the gradual recovery of the *ccs52a2-1* phenotypes, such as the SCN organization, possibly due to increased expression of *CCS52A1* in these tissues. At the shoot level, the CRISPR *pkn1 ccs52a2-1* double mutants show a partial leaf size recovery, owing to both an increased cell number and cell size, accompanied by a reacquisition of the wild-type DNA ploidy levels. Both the *pkn1-1 ccs52a2-1* and *pkn1-2 ccs52a2-1* mutants show a recovery in leaf size as well, albeit less than what was observed for the CRISPR *pkn1 ccs52a2-1* double mutants. In *pkn1-1 ccs52a2-1*, this recovery was caused by an increase in cell number and not cell size, again indicating that this mutant likely holds a secondary mutation that might be influencing leaf cell expansion without affecting endoreplication. This attests to the recently emerging evidence that although higher ploidy is often correlated with cell growth, it is itself not the major driving force for growth, but rather creates the circumstances in which growth through turgor-driven expansion can occur by inducing rapid cell wall changes (Bhosale et al., 2019). On the other hand, in *pkn1-2 ccs52a2-1*, the smaller leaf growth recovery phenotype compared to *ccs52a2-1* was not due to an increase in cell number, which was even further decreased compared to both the *ccs52a2-1* and *pkn1-2* single mutants, but instead was driven by a dramatic increase in cell size. Indeed, severe defects in cell proliferation have been observed to trigger a compensatory increase in cell size in determinate organs like leaves (Hisanaga et al., 2015).

### PKN1 as a Target of APC/C^CCS52A2^

Our data indicate that PKN1 is a direct target of the APC/C^CCS52A2^. Firstly, there is the fact that *pkn1* knock-out mutations can partially recover *ccs52a2-1* dwarf growth phenotypes. Secondly, the PKN1 amino acid sequence contains several degrons, forming putative recognition sites for APC/C^CCS52A2^, with their presence being conserved in PKN1 orthologs throughout the plant kingdom. Thirdly, PKN1 protein levels are increased upon treatment with a proteasome blocker, making it a likely target of ubiquitination-mediated proteasomal degradation. Finally, *ccs52a2-1* mutant plants with a PKN1 complementation or translational reporter construct show even more severe growth phenotypes, with roots even shorter and sporadically presenting doubled meristems or split root tips, indicating that the timely marking for breakdown of PKN1 by the CCS52A2-activated APC/C is essential for proper plant development. However, attempts made to prove direct interaction between PKN1 and CCS52A2 or other APC/C subunits remained fruitless, leaving the evidence for the control of CCS52A2 over PKN1 levels to be indirect.

### The Potential Role of PKN1 in the Cell Cycle

Several lines of evidence point towards PKN1 being a novel plant lineage-specific protein with a rate-limiting role during cell cycle progression. Not only was its expression correlated with cell division in both Arabidopsis and Marchantia, we demonstrated as well that the Arabidopsis PKN1 protein displays a cell cycle-dependent subcellular localization. Furthermore, Arabidopsis plants lacking functional PKN1 display a reduction in root meristem cell number, while those ectopically expressing *PKN1* show an increase in leaf cell number. Likewise, knock-out of the Marchantia *PKN1* gene resulted in a reduction in cell number. For Arabidopsis, it is curious that knock-out and overexpression of *PKN1* seem to specifically affect the root and the leaf, respectively, especially because the gene is expressed in both the root meristem and the developing leaf. One explanation might be that leaf cell number was analyzed at a timepoint where the leaf is thought to be mature (De Veylder et al., 2001; Kheibarshekan Asl et al., 2011). Loss of PKN1 likely only slows down the cell cycle, but does not block it, possibly leading only to a delay in leaf development, while still reaching the same end-point cell number as the wild type. In contrast to the leaf, the root meristem is constantly dividing to support continuous root growth, making a delay in cell division more readily visible as a decrease in the pool of meristematic cells present. The fact that plants overexpressing *PKN1* do not show any specific root phenotype might be explained by the fact that the *CaMV 35S* promoter used here can be less active in the root meristem than in other tissues (Ambrose et al., 2011), which was confirmed by the lower level of excess *PKN1* transcripts observed in the root compared with the leaf in these plants.

Arabidopsis PKN1 was observed to display a dynamic localization pattern during the cell cycle, being present at the preprophase band, the spindle and the phragmoplast during preprophase, mitosis and cytokinesis, respectively. Its potential to associate with microtubuli was confirmed by tobacco leaf infiltration experiments, where the *proPKN1:PKN1-GFP* signal could be detected at the cortical microtubules. Following cytokinesis, the reporter signal appeared to decline, which would fit the timing of APC/C^CCS52^ activation. A similar temporal degradation pattern in a CCS52A2-dependent manner was observed for the cell wall biosynthesis enzyme CSLD5, being important for cellulose deposition at the growing cell wall during cell division (Gu et al., 2016). The data suggest that some of the *ccs52a2-1* phenotypes might be attributed to altered microtubuli dynamics due to the inability to destroy PKN1. Such a role might be conserved, seeing the microtubule-associated localization pattern of the Marchantia PKN1 protein. Knock-out of *PKN1* in the *ccs52a2-1* mutant background might initially help to overcome such problems, but may eventually affect normal progression through mitosis as hinted by the cell division phenotypes in the *pkn1* mutants, probably explaining the rescue of the early but not the late developmental phenotypes of *ccs52a2-1*.

### PKN1 and CYCA3;4

Previously, the causal mutation for another revertant from the same EMS mutagenesis screen, *pkn2 ccs52a2-1*, was shown to be located in the gene coding for cyclin A3;4 (*CYCA3;4*) (Willems et al., 2020). The expression profiles of *PKN1* and *CYCA3;4* are markedly similar (Bulankova et al., 2013; Willems et al., 2020) and both genes appear to be co-expressed (Supplemental Fig. S7). Moreover, the PKN1 amino acid sequence holds several conserved potential CDK phosphorylation sites, making it a possible phosphorylation target of a CDK activated by CYCA3;4. However, we failed to prove a biochemical interaction between PKN1 and CYCA3;4. Moreover, the respective phenotypes of *PKN1* and *CYCA3;4* knock-out and overexpressing plants are different (Willems et al., 2020). Whereas *pkn1* mutant plants show a reduced root meristem cell division, plants lacking CYCA3;4 seem to have no growth phenotypes. Similarly, *PKN1* and *CYCA3;4* overexpressing plants show an increase and decrease in leaf cell number, respectively. Additionally, recovery of *ccs52a2-1* growth phenotypes in the *pkn1 ccs52a2-1* and *pkn2 ccs52a2-1* double mutants is not the same (Willems et al., 2020). Root growth in the days immediately after germination (3-5 DAS) is substantially better in the *pkn1 ccs52a2-1* mutants than in *pkn2 ccs52a2-1*, correlating with a more improved root tip tissue organization at 5 DAS. The opposite is seen in later development (6-9 DAS), correlating with *pkn2 ccs52a2-1* showing an almost full root meristem cell number recovery in contrast to the *pkn1 ccs52a2-1* mutants whose cell number is not markedly improved compared to *ccs52a2-1*. Moreover, although leaf phenotype recovery in the two revertants is similar and mostly due to an increased cell number, cell size recovery is markedly better in the CRISPR *pkn1 ccs52a2-1* double mutants than in *pkn2 ccs52a2-1*. Taken together, it seems most likely that PKN1 and CYCA3;4 perform independent functions controlling different aspects of meristematic cell division, while being similarly controlled at the protein level by APC/C^CCS52A2^-mediated breakdown.

In conclusion, PKN1 appears to be a novel and conserved plant-specific protein playing a role in cell division, likely through its interaction with microtubuli, and is proteolytically controlled by APC/C^CCS52A2^. To determine its exact role during mitosis and its influence on intracellular dynamics, future research might focus on identifying PKN1-interacting proteins, which will also enable the elucidation of the function of its different conserved amino acid regions and shed light on its protein structure.

## MATERIALS AND METHODS

### Plant Medium and Growth Conditions

*Arabidopsis thaliana* seeds were sterilized in 70% ethanol for 10-15 min and subsequently washed with 100% ethanol, after which they were left to dry in sterile conditions. For all experiments, the seeds were stratified in the dark for 2 days at 4°C before being placed in the growth room. Plants were grown *in vitro* under long-day conditions (16-h light/8-h dark, Lumilux Cool White lm, 50 to 70 µmol m^−2^ s^−1^) at 21°C on solidified half-strength Murashige and Skoog (MS) medium (2.151 g/L, 10 g/L sucrose, and 0.5 g/L 2-(N-morpholino) ethanesulfonic acid (MES), adjusted to pH 5.7 with 1 M KOH and 8 or 10 g/L plant agar). For analysis of root or shoot phenotypes, plants were grown vertically or horizontally, respectively. The treatments with MG132 were performed at a concentration of 100 µM for 24 h.

*Marchantia polymorpha* accession Takaragaike-1 (Tak-1) plants were grown under continuous light conditions at 21°C on solidified half-strength Gamborg’s B5 medium (Duchefa Biochemie) with 10 g/L plant agar.

### Constructs and Lines

The *ccs52a2-1* mutant line has been described previously (Vanstraelen et al., 2009), whereas the *pkn1-1 ccs52a2-1* double mutant was obtained through EMS mutagenesis of *ccs52a2-1* mutant seeds (see below). The *pkn1-1* single mutant was obtained from a segregating population resulting from backcrossing *pkn1-1 ccs52a2-1* with wild-type Col-0 followed by selfing and genotyping. The *pkn1-2* (SALK_047285) mutant was obtained from the Salk Institute T-DNA Express database (Alonso et al., 2003), and the *pkn1-3* to *pkn1-6* lines were obtained via CRISPR-based targeted mutagenesis in the Col-0 background (see dedicated paragraph).

The *proWOX5:GFP-GUS* transcriptional reporter was previously described (Heyman et al., 2016). The *PKN1* complementation construct *pFAST-R01-PKN1* was created by cloning a fragment containing the *PKN1* promoter (1500 bp directly upstream from start codon), gene sequence and 3’ untranslated region (300 bp directly downstream from stop codon) from Col-0 into the pDONR221 vector (Invitrogen) via a BP reaction and recombining it into the pFAST-R01 vector (Shimada et al., 2010) via an LR reaction. The *PKN1^OE^* construct was created by cloning the *PKN1* open reading frame (ORF) including stop codon from Col-0 into pDONR221 (Invitrogen, creating pDONR221-PKN1) via a BP reaction and subsequently recombining it via an LR reaction behind the strong *Cauliflower Mosaic Virus* (*CaMV*) 35S promoter in the pB7WG2 vector (Karimi et al., 2002). To create the PKN1 translational reporter lines, a PKN1 promoter fragment (consisting of 1500 bp upstream of the start codon) was cloned via a BP reaction into the pDONR-P4-P1r entry vector (Invitrogen, creating pDONR-L4-proPKN1-R1) and the *PKN1* ORF without stop codon was cloned via a BP reaction into pDONR221 (Invitrogen, creating pDONR221-PKN1nostop). The *pro35S:PKN1-GFP* construct was created by recombining pDONR221-PKN1nostop via an LR reaction behind the *CaMV 35S* promoter and in front of GFP in the pK7FWG2 vector (Karimi et al., 2002). The *pro35S:GFP-PKN1* construct was created by recombining pDONR221-PKN1 via an LR reaction behind the *CaMV 35S* promoter and GFP in the pK7WGF2 vector (Karimi et al., 2002). The *proPKN1:PKN1-GFP* and *proPKN1:PKN1-GUS* translational reporter constructs were created by recombining pDONR-L4-proPKN1-R1, pDONR221-PKN1nostop and pEN-R2-F*-L3 (GFP) or pEN-R2-S*-L3 (GUS) via an LR reaction into the pK7m34GW destination vector (Karimi et al., 2005; Karimi et al., 2007). All primer sequences used for cloning and genotyping are listed in Supplemental Table S2.

All vector-based cloning for Arabidopsis constructs was performed using the Gateway system (Invitrogen) according to the manufacturer’s recommendations. All constructs were transferred into the *Agrobacterium tumefaciens* C58C1RifR strain harboring the pMP90 plasmid. The obtained Agrobacterium strains were used to generate stably transformed Arabidopsis lines with the floral dip transformation method (Clough and Bent, 1998). All constructs were transformed into the Col-0 background, except for the *PKN1* complementation construct, which was transformed into *pkn1-1 ccs52a2-1*. Successful transformants were selected using kanamycin or Basta (glufosinate ammonium) or using fluorescence microscopy in the case of FAST constructs. In the T2 generation, only lines containing a single insert location of the construct were retained, while homozygous lines were selected in the T3 generation. Double mutants were made by crossing and confirmed through genotyping with PCR and/or sequencing.

For Marchantia constructs, the Tak-1 *MpPKN1* (Mapoly0057s0017, Mp7g06500) ORF without stop codon was cloned into pDONR221 (Invitrogen, creating pDONR221-MpPKN1nostop) via a BP reaction. The *pro35S:MpPKN1-GFP* construct was created by recombining pDONR221-MpPKN1nostop via an LR reaction behind the *CaMV 35S* promoter and in front of GFP in the pH7m34GW vector (Karimi et al., 2002). The *pro35S:MpPKN1-Tdtomato* construct was created by recombining pDONR221-MpPKN1nostop via an LR reaction into the pMpGWB130 vector (Ishizaki et al., 2015). Agrobacterium-mediated *M. polymorpha* Tak-1 transformation was done as previously described (Kubota et al., 2013). Transformants were selected on medium containing 10 mg/L hygromycin and 100 µg/ml cefotaxime.

### Generating CRISPR mutants

To generate the *A. thaliana PKN1* CRISPR mutants, two gRNAs specific for *PKN1* (gRNA1, TCAGTCTCAGCCACAGCCGC; gRNA3, ACTGATCTCACTGGTCTACG) were selected using the CRISPR-P web tool based on the expected cut site in the beginning of the gene (behind base pair 64 and 286 of the *PKN1* coding sequence, respectively) and minimal off-target scores (Lei et al., 2014). Primer pairs for the two gRNAs were designed to include 5’-end overhangs, being ATTG for the forward primer and AAAC for the reverse complement primer. The gRNA1 and gRNA3 primer pairs were inserted into the pGG-A-ATU6PTA-B and pGG-B-ATU6PTA-C vectors, respectively, after which these were recombined with the pGG-C-LinkerII-G vector into the pEN-L1-AG-L2 acceptor plasmid using Golden Gate, as previously described (Houbaert et al., 2018). This gRNA Gateway entry clone was then recombined with the pEN-L4-RPS5P-Cas9PTA-G7T-R1 and pB7m24GW3-FAST destination vectors in a Multisite Gateway LR reaction according to the manufacturer’s recommendations (Karimi et al., 2007; Houbaert et al., 2018). The construct was transformed into Col-0 plants as described in the previous paragraph, and successful transformants were selected using fluorescence microscopy on T1 generation seeds. T2 generation plants without the CRISPR construct were obtained by looking for seeds without fluorescence obtained from different T1 generation plants. Plants mutated at the expected sites were identified through SNP-genotyping PCR and sequencing, as previously described (Houbaert et al., 2018). All primer sequences used are listed in Supplemental Table S2.

To generate the *M. polymorpha PKN1* CRISPR mutants, the established CRISPR/Cas9-mediated mutagenesis system in *Marchantia* was used to generate *MpPKN1* mutant lines (Sauret-Güeto et al., 2020). Two different single gRNAs specific for *MpPKN1* (gRNA1: 5′-GAAAACCGACCAGCATGGTA-3′ and gRNA2: 5′-CTCGAAATCCGTAGCGCCTC-3′) were selected using the CRISPR direct web tool (https://crispr.dbcls.jp/) based on the expected cut site in the beginning of the gene (behind base pair 63 and 344 of the *PKN1* coding sequence, respectively) and minimal off-target scores. The gRNA1 and gRNA2 primer pairs were inserted into the L1_lacZgRNA-Ck2 (OP-075) and L1_lacZgRNA-Ck3 (OP-074) vectors, respectively, after which these were recombined with the L1_CsR-Ck1(OP-062) and L1_Cas9-Ck4 (OP-073) vector into the pCsA (OP-005) acceptor plasmid using Golden Gate Loop Assembly according to the manufacturer’s recommendations (Sauret-Güeto et al., 2020). The construct was transformed into Tak-1 background plants and transformants were selected with 0.5 μM chlorsulfuron (Sigma-Aldrich) and 100 µg/ml cefotaxime, fully frame-shifted transgenic lines were selected for further experiments. To genotype the mutant lines carrying a large deletion or inversion, the following primer sequences were used: 5′- TCACAACATACGTCCGACTAC -3′ and 5′- TCGGAGTGACGAGGGATGAA-3′.

### Plant Growth Phenotyping

Root growth and length were determined by marking the position of the root tip each day from 3 to 9 DAS, scanning the plates at 9 DAS and measuring using the ImageJ software package. Root meristem analysis was performed with the ImageJ software package using images of the root tip obtained with confocal microscopy, the distance between the QC and the end of the division zone was measured to determine the root meristem length, the number of cortical cells within the division zone was counted to determine the cortical cell number, and the average cortical cell size was determined by dividing the measured meristem length by the number of cortical cells.

To determine the rosette size, pictures were taken at 21 DAS using a digital camera fixed in position, after which the images were made binary (black and white) and the projected rosette size was measured using the wand tool in ImageJ. For analysis of leaf parameters, the first leaf pairs were harvested at 21 DAS and cleared overnight using 100% ethanol. Next, leaves were mounted on a slide with lactic acid. The total leaf area was determined from images taken with a digital camera mounted on a Stemi SV11 microscope (Zeiss) using ImageJ software. A DM LB microscope (Leica) with a drawing-tube attached was used to generate a pencil drawing of a group of at least 30 cells of the abaxial epidermis. On each leaf, the area chosen for drawing was located between 25 to 75% of the distance between the tip and the base of the leaf, halfway between the midrib and the leaf margin. After measuring the total drawn area (using the wand tool of ImageJ) and counting the number of pavement cells and stomata drawn, the average cell size, total number of cells per leaf and the SI (number of stomata divided by total number of epidermal cells) were calculated.

For *Marchantia* gemma and gemmaling size, the obtained confocal images were made binary (black and white) and the plant size was measured using the wand tool in ImageJ. For *Marchantia* gemma and gemmaling epidermal cell number, the obtained confocal images were analyzed in ImageJ running a macro script from The BioVoxxel Image Processing and Analysis Toolbox as previously described (Li et al., 2020). Epidermal cell density was calculated by dividing the plant size by the epidermal cell number.

### Flow Cytometry

For flow cytometry analysis, leaf material was chopped in 200 μL nuclei extraction buffer, after which 800 μL staining buffer was added (Cystain UV Precise P, Partec). The mix was filtered through a 30-μm green CellTrics filter (Sysmex – Partec) and analyzed by the Cyflow MB flow cytometer (Partec). The Cyflogic software was used for ploidy measurements. To calculate the EI, the following formula was used, with %nC representing the fraction of nuclei with n-times the haploid genome content: (0 x %2C + 1 x %4C + 2 x %8C + 3 x %16X + 4 x %32C) / Total nuclei

### Confocal Microscopy

For visualization of Arabidopsis root apical meristems, vertically grown plants were mounted in a 10-µM propidium iodide (PI) solution (Sigma) to stain the cell walls. Agroinfiltrated tobacco leaf cuttings were mounted in H_2_O. Imaging was done using an LSM 5 Exciter (Zeiss) confocal microscope. For PI and GFP excitation, the 543 line of a HeNe laser and the 488 line of an Argon laser were used, respectively. Laser light passed through an HFT 405/488/543/633 beamsplitter before reaching the sample, and emitted light from the sample first passed through an NFT 545 beamsplitter, after which it passed through a 650-nm long pass filter for PI detection, and through a 505- to 530-nm band pass filter for detection of GFP. PI and GFP were detected simultaneously with the line scanning mode of the microscope.

Marchantia plants were visualized using a LSM 710 (Zeiss) confocal microscope. Gemmae and 1-day-old gemmalings were mounted in a 10-µM PI solution (excitation 405 nm, emission 415–443 nm) as previously described (Westermann et al., 2020), while 2- and 3-day-old gemmalings were vacuum-infiltrated with ClearSee containing 0.1% calcofluor white (excitation 405 nm, emission 415–443 nm) prior to mounting according to the ClearSee protocol (Ursache et al., 2018). The *35S_pro_:MpPKN1-Tdtomato Marchantia* thalli were grown for 14 days in half-strength B5 medium before transfer to either DMSO control or oryzalin (10µM in DMSO) in half-strength B5 liquid medium for 1 h. To observe fluorescence, excitation at 561 nm and emission at 570-600 nm were used for TdTomato.

### EMS Mutagenesis

Roughly 14,000 *ccs52a2-1* seeds were subjected to EMS mutagenesis. The seeds were first hydrated with H_2_O for 8 h on a rotating wheel before being mutagenized with a 0.25% v/v solution of EMS for another 12 h. After treatment, seeds were washed twice with 15 mL 0.1 M sodium thiosulfate (Na_2_S_2_O_3_) for 15 min to stop the reaction and subsequently twice with H_2_O for 30 min. After that, seeds were left to dry on Whatman paper. Fifty-six pools of approximately 250 M_1_ seeds were mixed together with fine sand in Eppendorf tubes and sown in big pots with standard soil. After selfing, M_2_ seeds were harvested separately for every pool. Seeds were sterilized and sown on vertical plates to score for the reversion of the *ccs52a2-1* root growth phenotype. Plants with longer roots were subsequently selected and transferred to soil for self-fertilization. The recovery phenotype was then reconfirmed in the next generation (M_3_).

### Mapping of the Revertant Mutation

Segregating F2 progeny resulting from a cross between *pkn1-1 ccs52a2-1* and the *ccs52a2-1* parental line used for EMS mutagenesis was used as a mapping population. Approximately 250 plants showing the long root phenotype of the revertant were selected at 5 DAS and pooled for DNA extraction using the DNeasy Plant Mini Kit (Qiagen) according to the manufacturer’s instructions. DNA was extracted additionally from 200 plants of the original *ccs52a2-1* parental line. Illumina True-Seq libraries were generated from the extracted DNA according to the manufacturer’s protocol and sequenced on an Illumina HiSeq 100-bp paired-end run. The SHORE pipeline (Ossowski et al., 2008) was used for the alignment of sequences of both *pkn1-1 ccs52a2-1* and *ccs52a2-1* to the reference genome (Col-0; TAIR10). Using the SHOREmap pipeline (Sun and Schneeberger, 2015), an interval of increased mutant SNP alleles was identified and subsequently annotated. Filtering was performed within the interval for *de novo* EMS-specific SNPs with a concordance above 0.8 and intergenic or intronic mutations were removed to reveal the potential revertant mutations.

### SNP-genotyping PCR

The SNP-genotyping PCRs were performed as described previously (Mesrian Tanha et al., 2015), except that primer pairs specific for the wild-type or mutant allele were not combined in a single reaction, but rather used in separate PCR reactions in order to increase specificity. Prior to genotyping, test PCRs were performed to establish the most suitable annealing temperature for each used primer pair. All primer sequences used for the SNP-genotyping PCR are listed in Supplemental Table S2.

### Identification of homologs and conserved motifs

The dicotyledon PKN1 homolog sequences and that of *A. trichopoda* were obtained from Dicot PLAZA 4.5 (https://bioinformatics.psb.ugent.be/plaza/versions/plaza_v4_5_dicots/, (Van Bel et al., 2018)) under the homologous gene family HOM04D007290, while the monocotyledon homolog sequences were obtained from Monocot PLAZA 4.5 (https://bioinformatics.psb.ugent.be/plaza/versions/plaza_v4_5_monocots/, (Van Bel et al., 2018)) under the homologous gene family HOM04x5M006718. The gymnosperm PKN1 homolog sequences were obtained from Gymnosperm PLAZA 1.0 (https://bioinformatics.psb.ugent.be/plaza/versions/gymno-plaza/, (Proost et al., 2015)) under the homologous gene family HOM03D007930, which was identified using PLAZA’s internal BlastP functionality with default settings searching for sequences producing significant alignments with the *A.thaliana* PKN1 (alignment scores ranged from 47.8 to 60.5, with E-values ranging from 2e^-5^ to 1e^-9^). The PKN1 homologs of *M. polymorpha* and *P. patens* were identified using the internal BlastP functionality of Dicot PLAZA 4.5 with default settings, except that the E-value threshold was changed to 0.001, searching for sequences producing significant alignments with the *A.thaliana* PKN1 (alignment scores ranged from 50.4 to 52.8, with E-values ranging from 1e^-4^ to 3e^-5^). The PKN1 homologs of *S. moellendorffii* were identified using BlastP (https://blast.ncbi.nlm.nih.gov/Blast.cgi?PAGE=Proteins, (Johnson et al., 2008)) with the default settings, searching for sequences producing significant alignments with the *M. polymorpha* PKN1 within the *S. moellendorffii* non-redundant protein sequence database (alignment scores were 54.7, with an E-value of 3e^-7^), with sequences obtained from the RefSeq database (https://www.ncbi.nlm.nih.gov/refseq/). All sequences used are listed in Supplemental Dataset S1.

For motif discovery, the MEME (Bailey and Elkan, 1994) and MAST (Bailey and Gribskov, 1998) web-tools, part of the MEME Suite (https://meme-suite.org/meme/), were used. For input in MEME, a FASTA-type sequence file was compiled containing the PKN1 orthologs from the different plant species. When species contained multiple PKN1 paralogs, only the most conserved (i.e. most similar to *A. thaliana* PKN1) was used. Furthermore, homolog sequences that appeared to have been incorrectly annotated or were otherwise problematic, for example by being very short, split over multiple genes or missing stretches of sequence, were not used. See also Supplemental Dataset S1, column “Used for MEME”. MEME was run using the default settings, except that the number of motifs to be found was set at 10. The Supplemental Dataset S2 zip file contains the resulting MEME and MAST output web pages, available as HTML and as simple text file, as well as the exact sequences of the discovered motifs in the different species, available in FASTA and raw format. Logos of the ten different motifs were designed with the Weblogo3 web tool (weblogo.threeplusone.com/create.cgi), using the raw format motif sequences as input and using the default settings, with the following adjustments: Output Format was set at “PNG (high res)”, Sequence Type was set at “Protein”, Composition was set at “No adjustment for composition”, the Error Bars box was unchecked, the Version Fineprint box was unchecked, Y-axis Tic Spacing was set at “2”, and Color Scheme was set at “Chemistry (AA)”.

### RT-PCR and RT-qPCR

RNA was isolated with the RNeasy Mini kit (Qiagen) and was treated on-column with the RQ1 RNase-Free DNase (Promega) and used for cDNA synthesis with the iScript cDNA Synthesis Kit (Bio-Rad). For RT-PCR on the *pkn1-2* T-DNA line, cDNA made from RNA extracted from *pkn1-2* and Col-0 was used as a template for PCR using *PKN1* primers (see Supplemental Table S2) and the resulting amplicons were separated on a 1% agarose gel containing SYBRSafe (Invitrogen). RT-qPCR was performed using the SYBR Green kit (Roche) with 100 nM primers and 0.125 μL of RT reaction product in a total volume of 5 μL per reaction. Reactions were run and analyzed on the LightCycler 480 (Roche) according to the manufacturer’s instructions. For Arabidopsis, *EMB2386*, *PAC1*, and *RPS26E* were used as reference genes for normalization. For *Marchantia*, Mp*EF1α* (Mp3g23400, Mapoly0024s0116) and Mp*Actin* (Mp6g11010, Mapoly0016s0139) were used as reference genes for normalization (Saint-Marcoux et al., 2015). For each reaction, three technical repeats were performed. All primer sequences used for RT-qPCR are listed in Supplemental Table S2.

### GUS Staining

Plants were grown for the indicated period and fixed in an ice-cold 80% acetone solution for 3 h. Samples were washed three times with phosphate buffer (14 mM NaH_2_PO_4_ and 36 mM Na_2_HPO_4_) before being incubated in staining buffer (0.5 mg/mL 5-bromo-4-chloro-3-indolyl-β-D-glucuronic acid, 0.165 mg/mL potassium ferricyanide, 0.211 mg/mL potassium ferrocyanide, 0,585 mg/mL EDTA pH8, and 0,1% (v/v) Triton-X100, dissolved in phosphate buffer) at 37°C between 30 min and 16 h until sufficient staining was observed.

### Agroinfiltration

Wild-type tobacco (*N. benthamiana*) plants were grown under a normal light regime (14 h of light, 10 h of darkness) at 25°C and 70% relative humidity. The constructs were transferred using electroporation into the *A. tumefaciens* strain LBA4404 harboring the virulence plasmid VirG. The transformed Agrobacterium strain harboring the construct of interest was grown for 2 days in a shaking incubator (200 rpm) at 28°C in 10 mL of yeast extract broth medium, supplemented with the appropriate antibiotics. After incubation, the OD600 of each culture was measured with a spectrophotometer and an amount of culture representing an OD600 of 1.5 in a final volume of 2 mL was transferred to a fresh tube and spinned down. The resulting pellets were resuspended in 2 mL of infiltration buffer (10 mM MgCl_2_, 10 mM MES at pH 5.6, 100 µM acetosyringone) and 3x diluted with infiltration buffer (representing a final OD600 of 0.5) before infiltration. The inoculum was delivered to 3- to 4-week-old tobacco leaves by gentle pressure infiltration of the lower epidermis with a 1-mL syringe without needle. The infiltrated area of the leaf was delimited and labeled with permanent marker. The plant was incubated under normal growing conditions and expression of *GFP* was observed using confocal microscopy 3 to 5 days after infiltration. For pharmacological analysis, leaves infiltrated with *35S_pro_:MpPKN1-GFP* containing Agrobacterium were infiltrated after three days with DMSO or oryzalin (10 µM in DMSO) for a 1 h treatment, as described previously (Boruc et al., 2019).

### Accession Numbers

*A. thaliana* sequence data from this article can be found in the Arabidopsis Genome Initiative or GenBank/EMBL databases under the following accession numbers: *CCS52A2* (AT4G11920), *PKN1* (AT2G43990), *WOX5* (AT3G11260), *EMB2386* (AT1G02780), *PAC1* (AT3G22110) and *RPS26E* (AT3G56340). *Marchantia polymorpha* sequence data can be found in the PLAZA or MarpolBase databases under the following accession numbers: *MpPKN1* (Mapoly0057s0017 or Mp7g06500), Mp*EF1α* (Mapoly0024s0116 or Mp3g23400) and Mp*Actin* (Mapoly0016s0139 or Mp6g11010). The source and accession numbers of the *PKN1* orthologs can be found in Supplemental Dataset S1.

## SUPPLEMENTAL DATA

The following supplemental materials are available.

**Supplemental Figure S1.** Additional characteristics of the ccs52a2-1 and pkn1-1 ccs52a2-1 mutants.

**Supplemental Figure S2.** Detail of the allele frequency of EMS-specific mutations in *pkn1-1 ccs52a2-1*.

**Supplemental Figure S3.** Conserved motifs in the PKN1 protein.

**Supplemental Figure S4.** Analysis of the *pkn1-2* T-DNA line using RT-PCR.

**Supplemental Figure S5.** *PKN1* expression in wild-type shoots and roots.

**Supplemental Figure S6.** Intracellular localization of GFP-PKN1.

**Supplemental Figure S7.** Network of *PKN1* co-expressed genes.

**Supplemental Table S1.** Detailed annotation of the SNPs found for *pkn1-1 ccs52a2-1* in the interval selected on chromosome 2 from 17 Mbp up to the end.

**Supplemental Table S2.** Primer sequences.

**Supplemental Dataset S1.** *PKN1* homologs.

## ACKNOWLEDGMENTS

The authors thank Annick Bleys for help in preparing the manuscript.

